# Multi-scale transcriptome unveils spatial organisation and temporal dynamics of *Bacillus subtilis* biofilms

**DOI:** 10.1101/2023.01.06.522868

**Authors:** Yasmine Dergham, Dominique Le Coq, Pierre Nicolas, Julien Deschamps, Eugénie Huillet, Pilar Sanchez-Vizuete, Kassem Hamze, Romain Briandet

## Abstract

*Bacillus subtilis* has been extensively used to study the molecular mechanisms behind the development and dispersal of surface bacterial multicellular communities. Well-structured spatially organised communities (colony, pellicle, and submerged biofilm) share some similarities, but also display considerable differences at the structural, chemical and biological levels. To unveil the spatial transcriptional heterogeneity between the different communities, we analysed by RNA-seq nine spatio-physiological populations selected from planktonic and spatially organised communities. This led to a global landscape characterisation of gene expression profiles uncovering genes specifically expressed in each compartmental population. From this mesoscale analysis and using fluorescent transcriptional reporter fusions, 17 genes were selected and their patterns of expression reported at single cell scale with time-lapse confocal laser scanning microscopy (CLSM). Derived kymographs allowed to emphasise spectacular mosaic gene expression patterns within a biofilm. A special emphasis on oppositely regulated carbon metabolism genes (*gapA* and *gapB*) permitted to pinpoint the coexistence of spatially segregated bacteria under either glycolytic or gluconeogenic regime in a same biofilm population. Altogether, this study gives novel insights on the development and dispersal of *B. subtilis* surface-associated communities.

## INTRODUCTION

Spatially organised communities such as biofilms exhibit a set of microbial emerging properties and are embedded in a self-produced extracellular matrix ^1,2^. As these multicellular communities develop, bacteria adapt and respond differently to local chemical environmental conditions (*i*.*e*. concentration gradient of nutrient, oxygen, waste products and bacterial-signalling compounds), resulting in subpopulations of cells with considerable structural, physiological and biochemical heterogeneity over spatial and temporal scales^3^.

*Bacillus subtilis* has long served as a model organism for genetic studies on the formation of different types of surface communitie^4,5,6,7^. This Gram-positive, motile, spore-forming ubiquitous bacterium is frequently found in the rhizosphere in close proximity to plants, but also in extremely various environments^8,9^. It is commercially used to produce proteins, fermented food products, biocontrol agents and probiotic^10,11,12,13^. Conversely, it can potentially play a deleterious role, like the *B. subtilis* NDmed strain, isolated from a hospital endoscope washer-disinfector, capable of forming biofilms with complex protruding structures hyper-resistant to the action of oxidising agents used for endoscopes disinfection, thus protecting pathogenic bacteria such as *Staphylococcus aureus* in mixed-species biofilms^14,15,16^. Hence, understanding how these surface-bound communities are formed and interact is crucial for the development of suitable strategies for their control.

In an ever-changing environment, *B. subtilis* develops different adaptation strategies to survive including motility, matrix production and biofilm formation, sporulation, as well as induction of other stress responses^17,18,19^. In the laboratory, *B. subtilis* surface-associated multicellular community studies are typically based on the development of a floating biofilm or pellicle at the air-liquid interface, on a submerged biofilm at the solid-liquid interface, and on the development of complex colony at the solid-air interface^4,5,20^. In specific conditions, such as on a semi-solid surface, *B. subtilis* cells forming the colony can become highly motile and swarm over the surface by an organised collective movement while proliferating and consuming nutrients^21^. On a synthetic minimal medium, *B. subtilis* swarms from the multilayered colony in a branched, monolayer, dendritic pattern that continues to grow up to 1.5 cm from the swarm front. A transition from monolayer swarm to a multilayered biofilm occurs from the base of the dendrite and spreads outwards in response to environmental cues^22,23,24,25,26^. Thus, the *B. subtilis* NDmed strain has been phenotypically well characterised by multi-culturing approaches, which revealed its high ability to form 3D structures (colony, submerged and pellicle) and to swarm^4,6,27^.

A *B. subtilis* culture forming a biofilm contains at least seven different cell types: motile cells, surfactin producers, matrix producers, protease producers, cannibal, competent and sporulating cells^1,19,28,29^. This heterogeneity, which involves differential regulation of a number of genes, permits the division of labour between different cell types expressing different metabolic pathways^19,30,31,32,33,34^.

Temporal transcriptional analysis has been used to follow *B. subtilis* developmental strategies to form a complex biofilm. A study of metabolic changes during pellicle development by metabolomic, transcriptomic, and proteomic analysis, indicated that metabolic remodelling was largely controlled at the transcriptional level^35^. Besides, an ontogeny study of a *B. subtilis* macrocolony growing on agar has been shown to be correlated with evolution, and a temporal order of expression from older to newer genes^36^. Recently, we have performed a transcriptional study for the *B. subtilis* NDmed strain, for a whole static liquid model, in a microplate well, mixed and collected on a temporal scale^37^. This contributed to a first characterization of expression profiles during the first 7 hours of submerged biofilm development and for a mixture of different localised populations (submerged with detached cells and pellicle) after 24 hours of incubation^37^.

In the present study, we aimed to identify the differential expression of genes specifically expressed in different localised compartmental populations formed on solid, semi-solid or liquid interfaces. Hence, a spatial transcriptional analysis was performed at a mesoscopic scale for nine different localised multicellular populations, selected from planktonic culture, static liquid and swarming models. This has provided a global landscape characterization of gene expression for each of the differently selected populations. Comparison between the populations allowed to select 17 interesting genes whose expression was fluorescently reported for real-time monitoring using 3D and 4D imaging, paying special attention to single cell scale dynamics of the submerged biofilm population.

## RESULTS

### RNA sequencing shows spatially resolved compartmental populations with distinct patterns of gene expression

RNA-seq was used to compare transcriptomic profiles between multicellular localised populations from the different models formed by *B. subtilis* NDmed. All selected populations from the different compartments are schematized in Figure 1a. We have considered a 24-hour static liquid model where we have collected separately the submerged biofilm (SB), the floating pellicle (PL) and the free detached cells (DC). Moreover, from a swarming model, four differently localised compartments were collected; the mother colony (MC), the base of the dendrites (BS), the dendrites (DT) and the tips (TP). From the planktonic culture, the exponential (EX) and the stationary (ST) phases were collected, the latter being used as the inoculum to initiate the two models.

**Figure 1:**
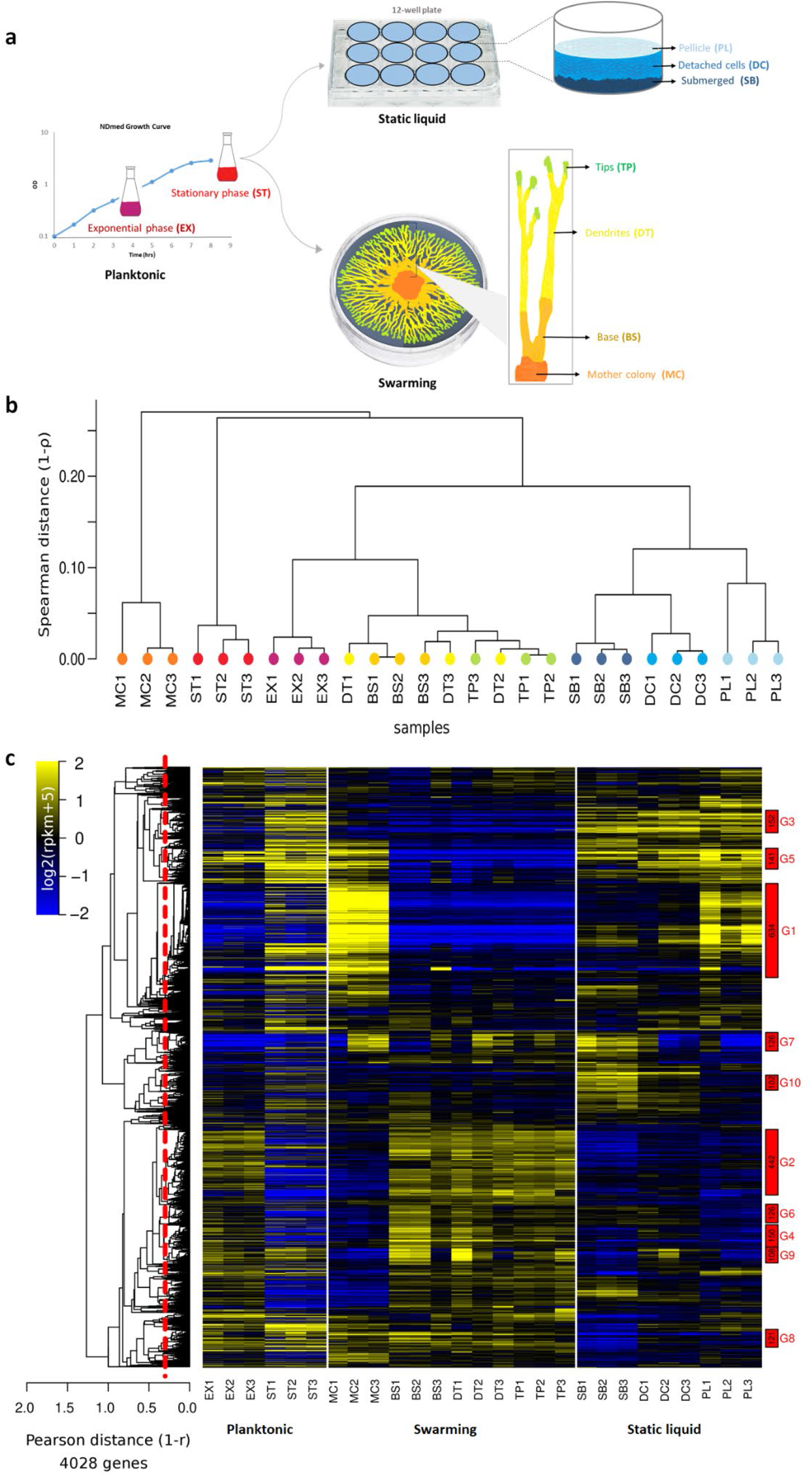
An overview of the spatial transcriptome remodelling between the different selected compartments of B. subtilis. (a) Schematic drawing of the differently localised spatial compartments selected. From the planktonic culture, the exponential (EX) and the stationary (ST) phase were selected. From the static liquid model, the pellicle (PL) formed at the liquid-air interface, the submerged biofilm (SB) formed on the solid-liquid interface, and the free detached cells (DC) between these two compartments were collected separately. From the swarming model, four localised compartments were collected separately: the mother colony (MC), the inoculation site from which the swarm has developed as a mature macrocolony; the base (BS) of the dendrites as an earlier biofilm form; the dendrites (DT), a monolayer of cells ready to form later the biofilm; and the tips (TP) formed of motile and highly dividing cells. For each compartment, three independent samples were taken as biological replicates. (b) Pairwise distance (Spearman) between RNA-seq profiles is summarised by a hierarchical clustering tree emphasising the divergence of the mother colony (MC) and the stationary phase (ST) between/along other selected compartments and the closeness of the adjacent spatial compartments of either the static liquid model (SB, DC and PL) or the swarming model (BS, DT and TP), which share a closer genetic expression profile with the exponential phase (EX). (c) Global heatmap representation for the 4028 genes present in NDmed across the spatially selected surface-associated compartments. The colour code reflects the comparison to the mean computed for each gene (log2 ratio) taking as a reference the average of all conditions, except the planktonic ones (EX and ST). The hierarchical clustering tree shown on the left side of the heatmap (average link) was cut at average Pearson correlation of 0.7 (dashed red line) to define the expression clusters shown as rectangles on the right side of the heatmap. Clusters were named (from G1 to G321) by decreasing sizes and only those containing more than 100 genes are highlighted (number of genes printed in black, cluster name printed in red).

To assess the quality and reproducibility of the RNA-seq data, a hierarchical clustering of the samples was performed (Fig. 1b). This analysis shows distinct transcriptomic profiles between the different spatial populations of compartments. The three biological replicates are grouped together. The only exception is for the adjacent swarming compartments (BS, DT and TP), where the clustering does not strictly group the samples by compartment but rather exhibits a trend to separate BS-DT from DT-TP samples. This could be due to the technical difficulty to precisely delineate visually these adjacent compartments, and/or because the physiology of the cells in the DT could be very similar either to that in the BS or in the TP. To investigate these global differences, a statistical analysis was conducted to identify differentially expressed genes (DEGs) between the compartments of each model (Supplementary Fig.S1). In line with the difficulty to reliably distinguish these three compartments (BS, DT, and TP), a pairwise comparison for these adjacent compartments identified 12 DEGs when comparing the DT to the BS and 24 DEGs when comparing the TP to the DT, with a strict increase in their number to reach 304 DEGs when comparing the TP to the BS (Supplementary Fig.S1).

Expression profiles for the 4028 genes of *B. subtilis* NDmed along all the considered conditions is presented in the heatmap Figure 1c. Groups of genes with a similar expression profile across samples were identified with a cut-off on average pairwise Pearson correlation within a group (r=0.7). The function of the genes within each of these groups of co-regulation were characterised using *SubtiWiki*-derived functional categories^38^ (Supplementary Fig.S2). The largest group (G1, 634 genes), upregulated in the MC and the PL, governs around 76% of the genes related to sporulation (Fig.1c, Supplementary Fig.S2). The second largest group (G2, 442 genes) contains 25% of the genes required for protein synthesis, modification and degradation, are upregulated in the EX and in the swarming compartments (BS/DT/TP). Around ∼68% of the genes involved in motility and chemotaxis are clustered in G9; they are downregulated in the three biofilm populations (MC, PL, and SB) and upregulated in all the other compartments. Genes required for biofilm formation are mainly found in G8 with ∼42% of them clearly downregulated in the SB (Fig. 1c). Unknown or poorly characterised putative genes constitute ∼39% of the genome, among which genes with interesting profiles, *i*.*e. ywdK, yodT, yjfA*, and *yezF* are highly expressed in the biofilm populations (MC, SB and PL); *yqbR, yqzN, yrhG, ydgG* highly expressed in the TP; *ykuO, ywmE, yfmQ* highly expressed in the PL; *yitJ, yhfS, yoaC, yaoD, yhfT, yxKC* strongly downregulated in the SB compared to other compartments. To better compare the population genetic expression levels between adjacent compartments and to highlight the different functional categories encoded by the differentially expressed genes on a spatial level, the static liquid and the swarming models were analysed separately in supplementary Fig.S3 and Fig.S4.

### Spatio-temporal patterns of gene expression reveals the various heterogeneous subpopulations present during biofilm development

Subjecting the whole compartment of a biofilm population to transcriptome analysis allowed us to assess the average gene expression genome-wide but we were interested in going beyond this mesoscale analysis by visualising gene expression *in situ* at a single cell level. For this purpose, based on the transcriptome data and known gene functions, we identified genes representing the different cell types present in a biofilm (scattered genes in the global heatmap, Supplementary Fig.S5), and constructed transcriptional fusions to fluorescent reporter genes *gfp* or *mCherry*. Matrix genes are represented by *epsA, tapA, bslA, srfAA, ypqP* and *capE*, motility by *hag*, exoprotease by *aprE*, carbon metabolism by *ackA, cggR, gapB*, competence by *comGA*, cannibalism by *skfA*, respiration by *ctaA* and *narG*, and sporulation by *spoIIGA* and *spoVC*.

Quantitative data from the transcriptome analysis were validated by an *in-situ* 3D microscopic observation in both swarming and static liquid models, using the different reported genes (Supplementary Fig.S6). Confocal imaging also pointed out spatial heterogeneity patterns of gene expression along the different selected compartments. Most of the reported genes show a lower or only moderately higher expression in the SB compared to the MC or the PL after 24 hours of incubation at 30°C (Supplementary Fig.S6). This suggests that these genes are either always weakly expressed in the SB, or expressed during a short window of time before or after our observation time-point (24 hours). This second hypothesis led us to monitor temporally the reported genes from 0 to 48 hours of incubation.

A real-time movie of Gfp expression by the NDmed-GFP strain at the submerged level, illustrated by a kymograph in Figure 2a, shows how cells adhere to the surface during the first few hours, then stop separating out and form sessile chains, followed by a sudden differentiation of a subpopulation into motile cells (between 5 and 10 hours of incubation). Only in a second kinetics sessile cells colonise the surface to form the highly structured SB (Supplementary Movie S1).

**Figure 2:**
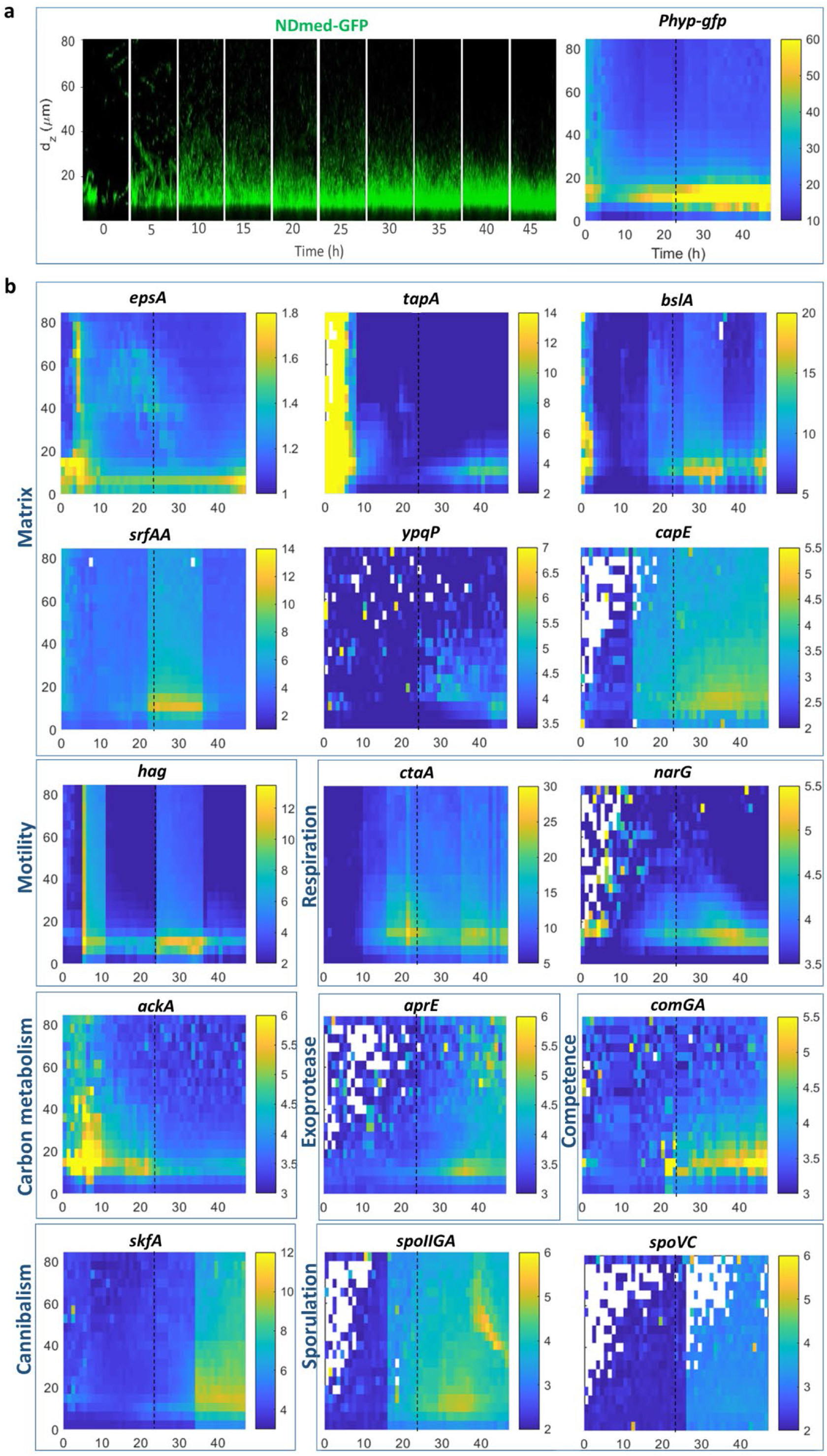
Spatio-temporal monitoring of gene expression in submerged biofilm (SB). (a) On the left is presented 4D confocal imaging (x 50μm, y 50μm, z 80μm) for the NDmed-GFP strain. A kymograph showing by a colour code the intensity of Gfp expression as a function of time and space is presented on the right. (b) Kymographs representing spatio-temporal expression of 15 transcriptional reporter fusions to genes potentially involved in biofilm development. The black dotted line in each kymograph represents the time (24 hour) corresponding to the RNA-seq analysis.

For all the reported genes, representatives of the main functional activities potentially present during biofilm formation, a temporal scale monitoring the intensity of gene expression is represented as kymographs in Figure 2b. Expression of *epsA* and *tapA*, involved in the synthesis of the major matrix components in a biofilm, is high during the first 5 hours of SB formation, followed by a global gradual decrease with some clusters remaining at high expression. Only after 30 hours, a slight increase of expression is observed, homogeneously scattered on the submerged level (Fig. 2b, Supplementary Movie S2). BslA, another structural protein in the biofilm matrix, acts synergistically with both TasA and EPS^32^. In the SB, expression of *bslA* is upregulated in a few clusters during the first 4 hours, followed by homogenization of a basal level of expression which increases progressively with time. The *ypqP* gene, involved potentially in the synthesis of polysaccharides participating in the strong spatial organisation^27^, shows some stochastic expression by very few cells at the beginning of biofilm formation; after 30 hours *ypqP* is expressed at a low level. In a similar manner expression of *capE*, involved in capsular polyglutamate synthesis, remains at a very low level between 11 to 25 hours of incubation to increase moderately afterwards. The *srfAA* gene, involved in surfactin synthesis, is weakly expressed for the first 18 hours and then strongly expressed in a time frame between 21 and 36 hours of incubation, to be downregulated afterwards. A burst of expression of *hag*, encoding flagellin, occurs after 5 hours of incubation, synchronised with the beginning of down-regulation of *tapA* (Fig. 2b, Supplementary Movie S2). A gradual decrease is then observed, followed by another wave of high expression of *hag* between 24 and 36 hours of incubation (Fig. 2b).

The *ctaA* gene, encoding a heme A synthase, is one of several genes involved in aerobic respiration regulated by ResD^39^. A significant expression is observed after 8 hours of incubation, followed by oscillations of high expression. Stochastic expression of anaerobic genes, represented by *narG*, is observed in very few cells during the first 5 hours of incubation, followed by a continuous gradual expression starting at around 14 hours.

For carbon metabolism, *ackA*, encoding acetate kinase, shows an upregulation during the first 12 hours, and is progressively downregulated after. This downregulation is faced by an upregulation of *aprE*, encoding the major extracellular alkaline protease (Fig. 2b, Supplementary Movie S3). A brutal expression of *comGA*, involved in competence acquisition, is seen in countable cells after 21 hours giving the high expression as appearing on the kymograph (Fig. 2b). This is then accompanied by an increase of the subpopulation expressing moderately *comGA* (Supplementary Movie S4). Expression of *skfA*, encoding the spore killing factor, is significantly observed from 21 hours with a noticeable increase in intensity after 31 hours of incubation (Supplementary Movie S4). We have also monitored the expression of *spoIIGA* and *spoVC* involved respectively in early and late sporulation steps. Figure 2b, shows that *spoIIGA* starts to be expressed at around 18 hours of incubation, indicating the beginning of sporulation, while expression of *spoVC* mainly starts after around 28 hours of incubation.

### Spatial transcriptome detects oppositely regulated subpopulations occurring side by side within a biofilm

Glycolysis and gluconeogenesis are two opposite pathways, for which *B. subtilis* possesses two distinct glyceraldehyde-3-phosphate dehydrogenases (GAPDH) (EC 1.2.1.12) catalysing either the oxidative phosphorylation of glyceraldehyde-3-phosphate into 1,3-diphosphoglycerate or the reverse reaction: (i) GapA, a strictly NAD-dependent GAPDH involved in glycolysis, and (ii) GapB, involved in gluconeogenesis and exhibiting a cofactor specificity for NADP^40^. Since coexistence of the two pathways in the same cell dissipate energy in a futile cycle^41^, expression of *gapA* and *gapB* are subjected to very efficient opposite regulations: *gapA* transcription is induced in glycolytic conditions and is repressed during gluconeogenesis by the self-regulated CggR repressor of the *cggR-gapA* operon, whereas *gapB* is transcribed only during gluconeogenesis and strongly repressed under glycolytic conditions by the CcpN repressor, as is *pckA*, encoding the purely gluconeogenic PEP-carboxykinase (PEP-CK)^40,42,43^.

Although cultures for this study were performed in purely glycolytic conditions (*i*.*e*. with glucose as a carbon source), we have observed (Fig. 3a) that in the three biofilm populations (MC, SB, and PL) expression of *gapB* and *pckA* was derepressed after 24 hours of incubation, indicating a depletion in glucose in these compartments. Interestingly, the two strictly oppositely regulated groups of genes, *cggR-gapA* on one hand, or *gapB* and *pckA* on the other hand, are oppositely regulated in all the different selected compartments, except in the SB where these 2 groups are both upregulated (Fig. 3a). This observation suggested coexistence of two cell types in the same compartment and motivated the construction of a strain reporting the expression of both *cggR-gapA* and *gapB* by different fluorescent transcriptional fusions (Table 1).

**Figure 3:**
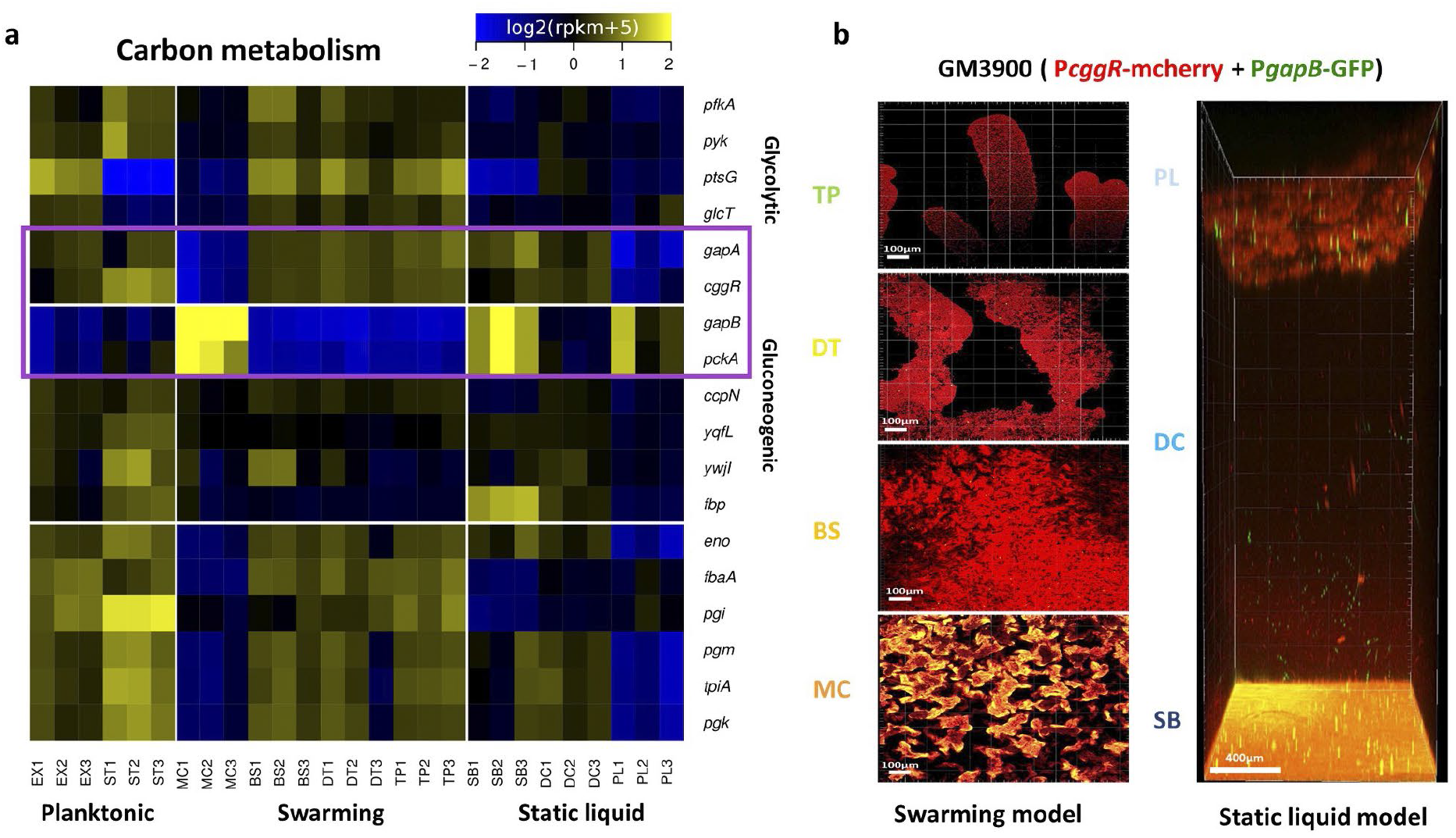
Spatial transcriptomic remodelling with in situ 3D imaging highlights a heterogeneous differential expression of central carbon metabolism. (a) Heatmap representation of the relative variations of expression level across samples. The colour code reflects the comparison to the mean computed for each gene across all the samples, except the planktonic (EX and ST) (log2 ratio). Genes were selected from Subtiwiki categories specific for glycolysis or gluconeogenesis, or common to both pathways (level 3). The purple box highlights central genes specific respectively for glycolysis (gapA, cggR) or gluconeogenesis (gapB, pckA). (b) Spatial confocal imaging for the different selected compartments from the swarming (MC, BS, DT, TP) and the static liquid (SB, DC, TP) models after 24 hours at 30°C. Using strain GM3900 reporting transcription of cggR-gapA by mCherry (in red) and of gapB by gfp (in green), with the same protocol as for the transcriptome analysis, except for the static liquid model the usage of 96-well microplate instead of the 12-well. Three replicative observations were performed independently for each model.

**Table 1.**
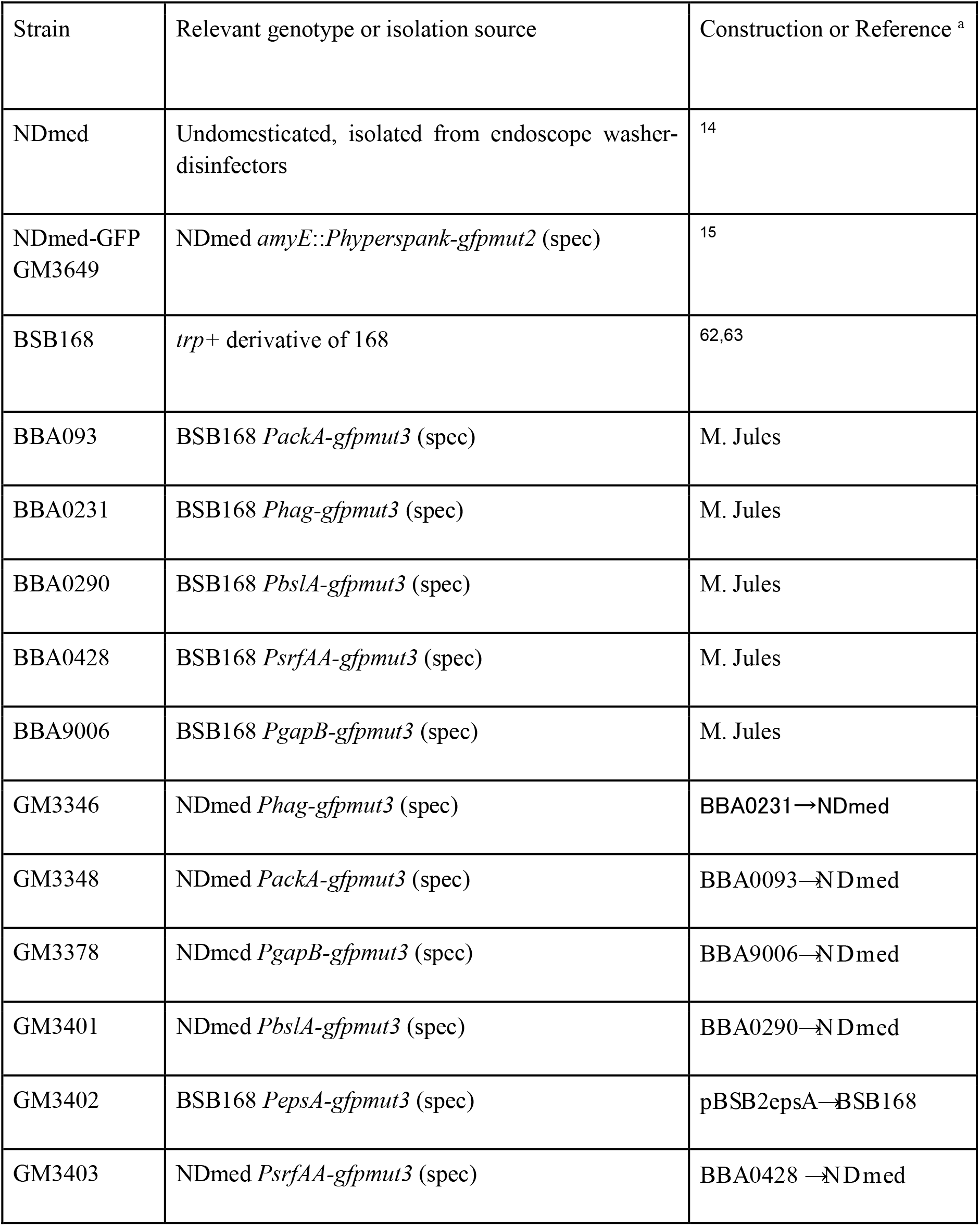
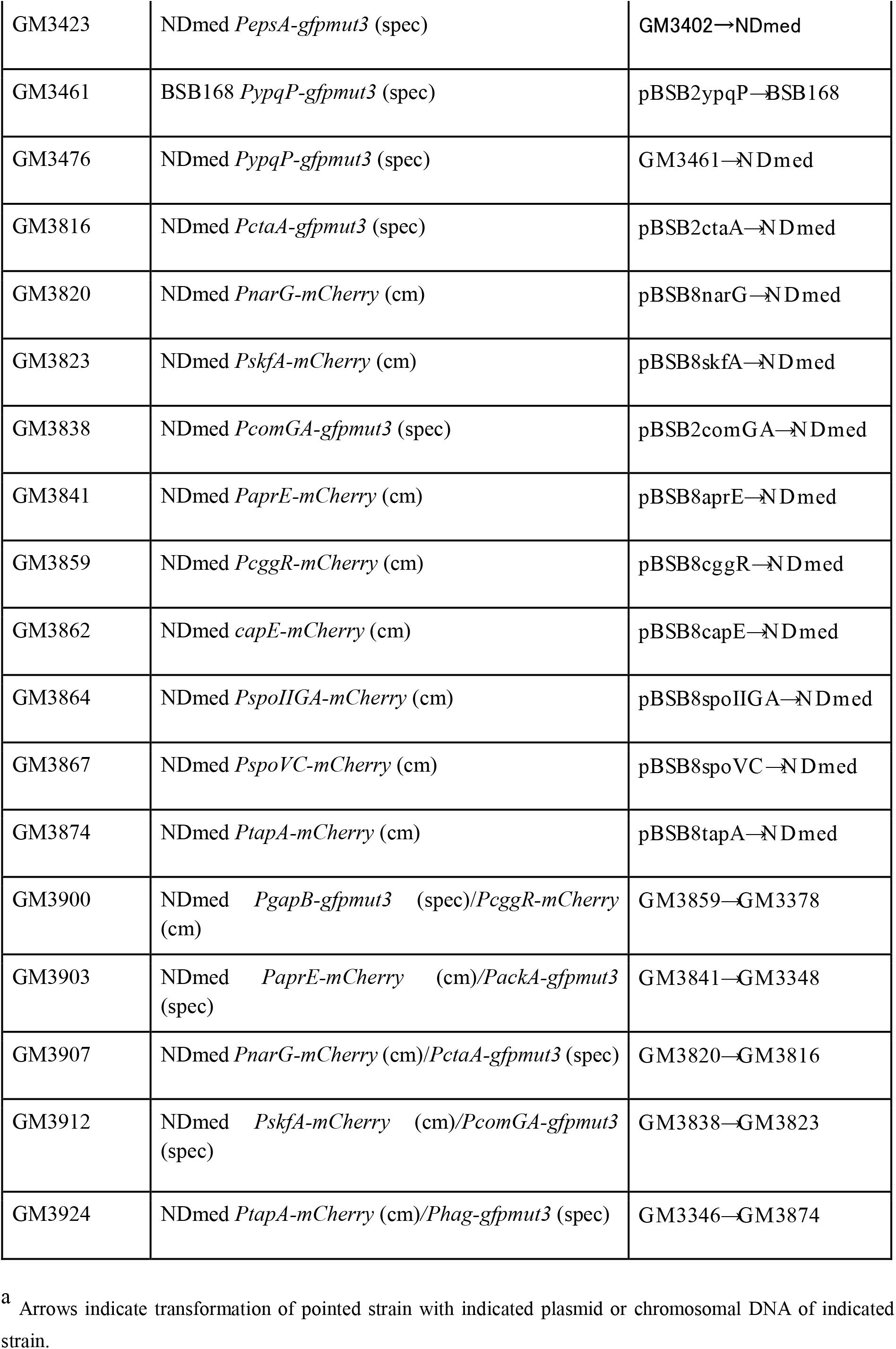
*B. subtilis* strains used in this study.

Using this strain (GM3900) and CLSM imaging, we observed at a single cell level the *in-situ* expression of both GapA and GapB in the different spatially localised populations of compartments. Figure 3b represents a real-time spatial monitoring after 24h for the different compartments of a swarming model (MC, BS, DT, and TP) and a static liquid model (PL, DC, and SB). Observation of GM3900 swarming on a glycolytic medium clearly confirmed our previous transcriptome data, from which the glycolytic genes *gapA* and *cggR* appeared upregulated all along the swarming compartments (BS, DT, and TP) and were rather downregulated in the MC. On the contrary, the gluconeogenic genes *gapB* and *pckA* were repressed in the swarming compartments, and highly upregulated in the MC. In the static liquid model, gluconeogenic genes were upregulated in both the PL and the SB compartments; an upregulation of glycolytic genes was also observed in both the SB and the DC compartments. Microscopy observations allowed to display the coexistence of subpopulations under either a glycolytic or a gluconeogenic metabolic regime in all the three biofilm compartmental populations (PL, SB and MC).

### Conversion from glycolytic to gluconeogenic regime starts from localised single cell within a glycolytic expressing population

Most of the physiological and genetic studies on regulation of carbon central metabolism and glycolysis/gluconeogenesis have been performed with planktonic liquid cultures in defined media, laboratory conditions not reflecting the complexity of the regulations involved in bacterial natural habitats. A closer observation to the BS, in the swarming model, or to the PL, in the static liquid model, allowed to visualise in GM3900 the switch to a gluconeogenic regime occuring in a single cell from a population growing with a glycolytic metabolism (Fig. 3, Supplementary Fig.S7). We then performed *in situ* spatio-temporal scale monitoring for the submerged compartment with a higher resolution (Fig. 4). 4D confocal imaging of GM3900 shows a high expression of glycolytic genes in bacteria growing in the glycolytic B medium (Supplementary Movie S5). After 15 hours of incubation, as nutrients become limited, there is a gradual decrease of cells expressing glycolytic genes, followed by a sudden expression of gluconeogenic genes in small clusters of few cells. Cells under a glycolytic regime continue to decrease with an upregulation of gluconeogenesis in a few other cells. Then after 22 hours cells regain a glycolytic metabolism and after 24 hours most of the population is again in glycolysis, but with some clusters of cells in gluconeogenesis (Fig. 4). After 24 hours, one can observe a slight increase in the number of cells of both subpopulations expressing either glycolytic or gluconeogenic genes being spatially mixed together, followed after 42 hours by strict increase in subpopulations expressing opposite carbon metabolism regulatory pathways (Supplementary Movie S5). Even after prolonged incubation these subpopulations seem to remain associated together. These observations indicating the coexistence of spatially mixed subpopulations growing either under a glycolytic or gluconeogenic regime in all the three biofilm populations (MC, PL and SB) suggest the existence of metabolite exchange between these subpopulations. Besides, another source of metabolites could be provided by dead cells.

**Figure 4:**
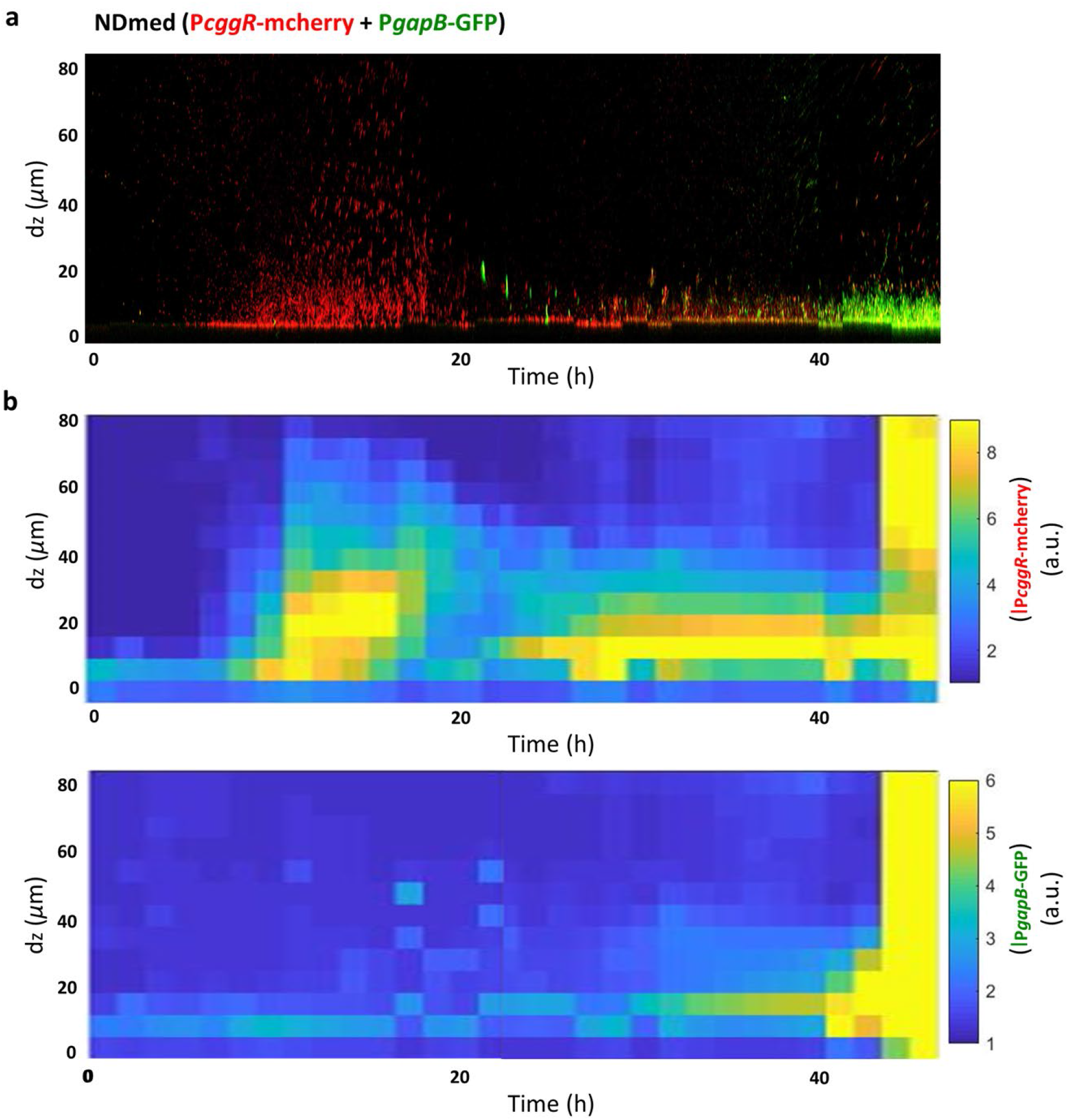
Spatio-temporal imaging for the submerged biofilm compartment. **(a)** Sections from a real-time confocal imaging (x 50μm, y 50μm, z 80μm) for 48 hours, image every one and half hour, using strain GM3900 reporting transcription of cggR (gapA) by mCherry (in red) and of gapB by gfp (in green), with the same protocol used for the transcriptome analysis, except the usage of 96-well microplates instead of the 12-well (Supplementary Movie S5). **(b)** Kymographs representing by a colour code the intensity of the expression for the transcriptional reporter fusions to the cggR and gapB genes along a spatio-temporal scale. Three biological replicates were performed.

### Two successive waves of localised cell death remodel the biofilm organisation

To further understand the heterogeneity and fluctuations of the different functions during *B. subtilis* biofilm development, a Live/Dead tracking at single cell scale was performed. Figure 5a, represents kinetic images for the live cells of *B. subtilis* reported by their expression of green fluorescent protein (GFP, green), while the dead cells and eDNA were contrasted with propidium iodide staining (PI, red). A multidimensional kymograph representing the intensity of dead cells (obtained by a ratio of dead/live cells) as a function of their spatial localization and time is presented in Figure 5b. Bacteria adhere to the surface and form chains of sessile cells in the first few hours of incubation and thereafter, between 15 and 24 hours, clusters of dead cells are observed over the formed biofilm (Fig. 5, Supplementary movie S1). After this first wave, the dead cells density decreases (Fig. 5), faced by a slight increase in the live population until around 42 hours where a second wave of dead cells occurs (Fig. 5a, Supplementary movie S1). Interestingly, by comparing the kymographs in Figures 2a and 5b, it appears that these dead cells subpopulations are mainly spatially localised as a layer on the top of the SB live cells.

**Figure 5:**
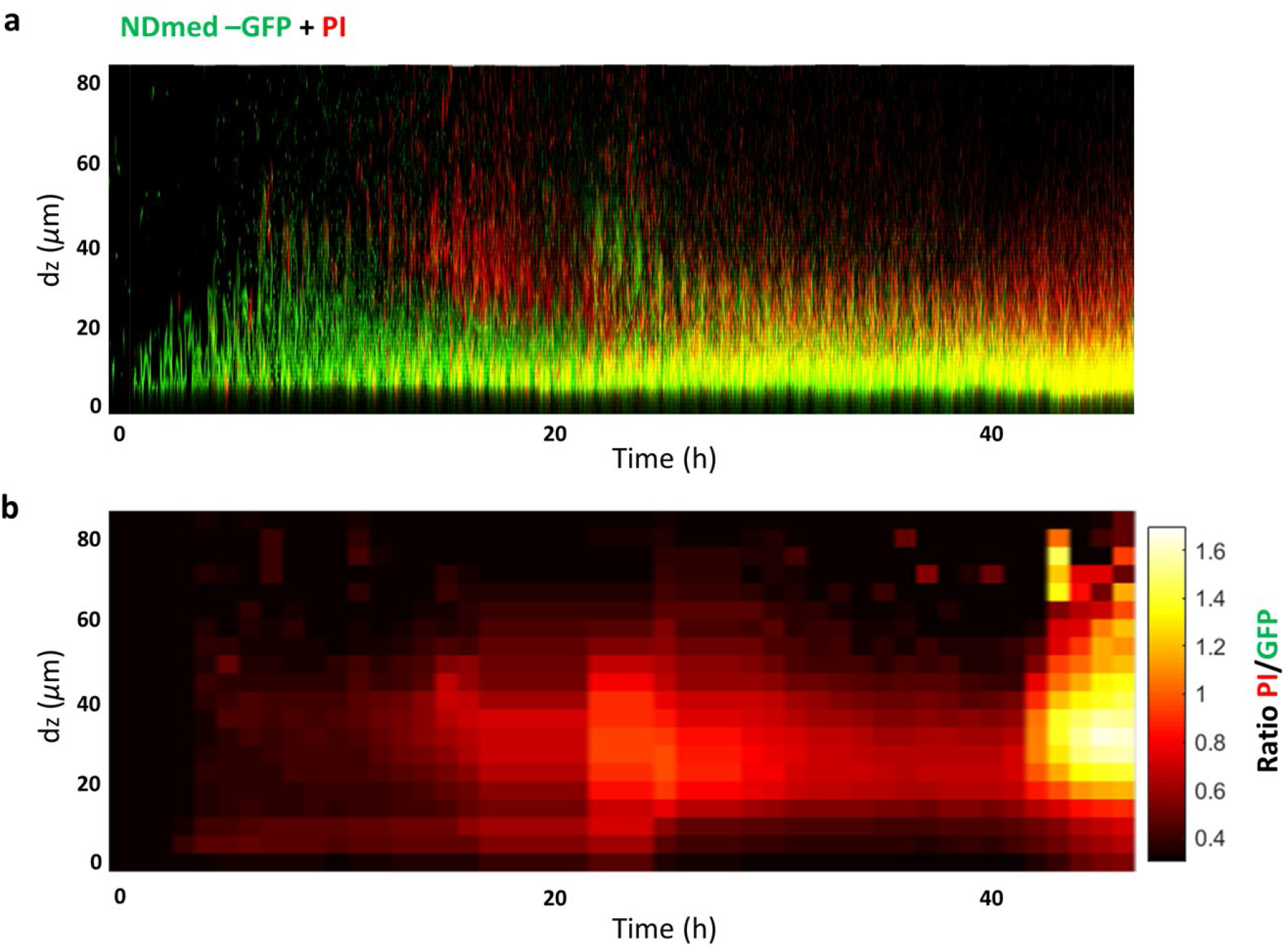
Temporal observation for the submerged biofilm (SB) development reveals oscillations of dead cell spatial localization. (a) Sections from a real-time confocal imaging (x 50μm, y 20μm, z 80μm), image every one hour, using NDmed-GFP (GM3649) and PI for permeable cell staining, with the same protocol used for the transcriptome analysis, except the usage of 96-well microplates instead of the 12-well (Supplementary Movie S1). (b) Kymograph representative of at least three replicates, representing the ratio of dead/live cells along a spatio-temporal scale.

**Figure 6:**
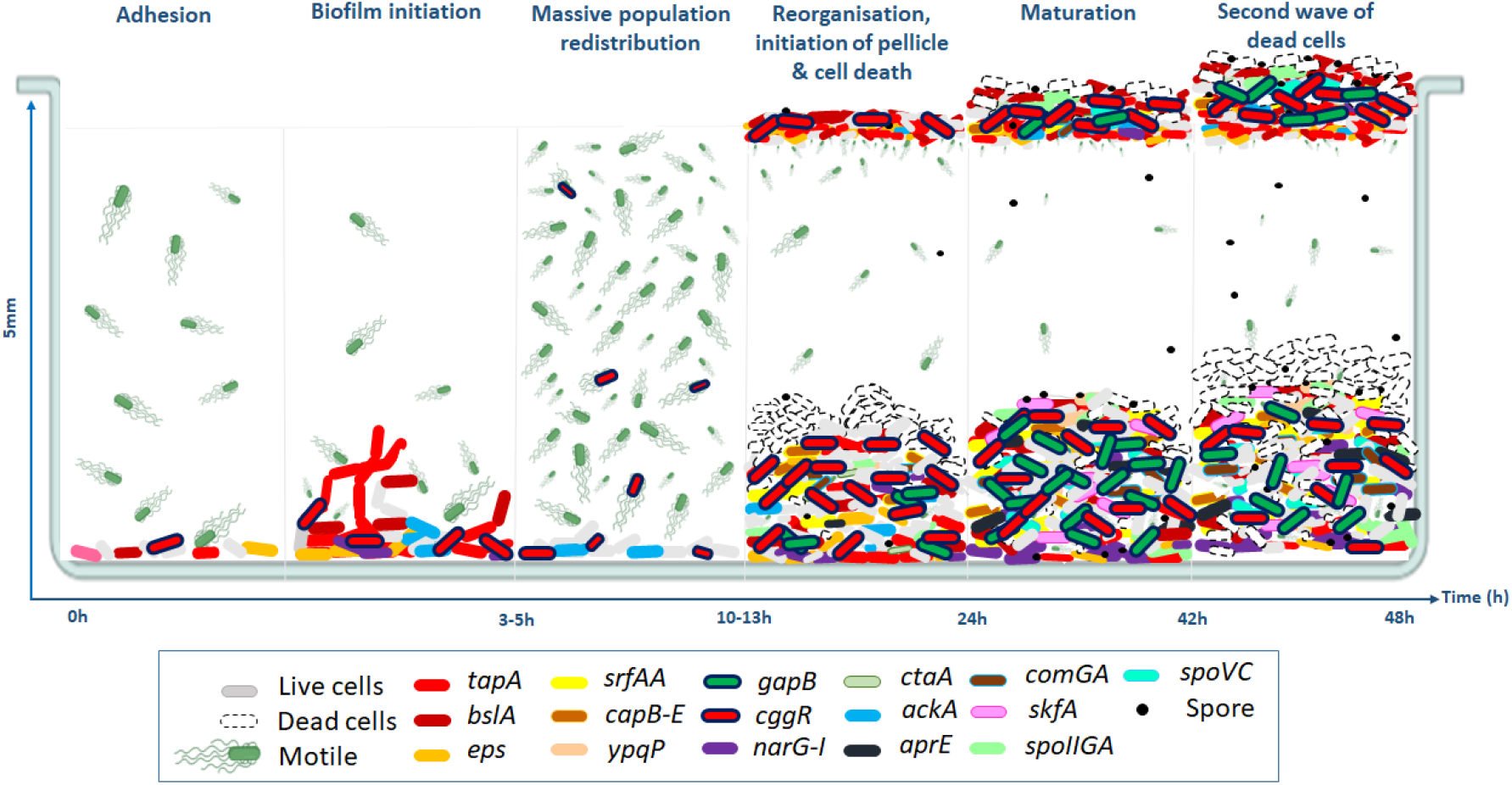
Spatio-temporal diversification of B. subtilis cell types in a well of microplate. A schematic illustration proposed for the static liquid biofilm dynamics over 48 hours, using a microplate and different reporting techniques. First, cells adhere to the submerged surface, followed by biofilm initiation where adherent sessile cells proliferate expressing matrix genes (i.e. tapA and eps). This is followed by a massive population redistribution, during which a sudden cell differentiation from sessile to motile cells occurs within a 15 minutes range. Then the submerged biofilm is reorganised and the formation of a pellicle is initiated at the air-liquid interface. This is accompanied by a 1^st^ localised cell death wave (between 13 and 24 hours). Maturation of biofilm, associated with a slight increase in the live population is followed by a 2^nd^ wave of cell death (after around 42 hours).

## DISCUSSION

Spatial transcriptomic data generated in this study put forward a global view on the variation of gene expression profiles for nine localised compartments, including three biofilm populations: the MC, PL and SB. A global hierarchical clustering of the RNAseq analysis (Fig. 1b) points out that the MC formed on agar showed a very distinct transcriptome profile compared to the PL and the SB. With the different environmental conditions, similarities could still be exhibited between the different biofilm populations. For instance, the *hag* gene encoding flagellin and reporting motility is downregulated in the three biofilms compared to the other compartments explored (DC, BS, DT, and TP). Microscopy observations of fluorescent transcriptional fusion further allowed to contrast minor subpopulations of cell expressing motility genes, in particular on the interfacial layers of the community, *i*.*e*. embedded under matrix-producing cells. This corresponds to the layer near to the agar surface for the MC; the inner immersed layer for the floating PL, and the layer in contact with the substratum for the SB (Supplementary Fig.S6). Expression of flagella could also be present within the biofilm indicating the migration of cells by chemotaxis toward a zone richer in oxygen and nutrients, allowing the vascularization of the biofilm matrix to increase diffusion/reaction throughout the biofilm^44^.

Most of the genes involved in sporulation appear strongly upregulated in aerial biofilms (MC and PL) and poorly expressed in the SB community. Surprisingly, in the static liquid model, a specific counting of spores indicated a higher quantity in the SB fraction than in the PL (data not shown), although sporulation genes were more expressed in the PL. Indeed, it has been shown recently that the spore surface of *B. subtilis* was covered with legionaminic acid, required for the crust assembly and enhancing hydrophilicity^45^. All together these observations suggest that, in the timeframe we explored, spores essentially produced in the PL can sediment as hydrophilic colloids down the well and accumulate within the SB level. Another striking difference between these compartments is the dominant anaerobic respiration metabolism detected in the SB compared to the other aerial biofilm populations. Within a static liquid model, the coexistence of two interfacial biofilm communities of *B. subtilis* with distinct respiration metabolisms is pointed out here: the SB (and the DC) mainly under anaerobic, and the PL under aerobic respiration. Although the PL and the MC are in contact with the air, the existence of a small subpopulation of cells expressing anaerobic genes is still observed. Taking advantage of RNA-seq data and of transcriptional reporter fusions associated with the microscopy technique, we could observe that the major extracellular matrix genes (*i*.*e. epsA-O, tapA* operons, and *bslA*) are more strongly and homogeneously expressed in the aerial biofilms than in the SB, which forms small dispersed clusters. This diversity in the spatial repartition of cells producing each of these matrix components suggests different biochemical matrix composition associated with specific local micro-rheological properties. For better visualisation of the genetic expression level among adjacent compartments, the swarming model and the static liquid model were analysed separately (Supplementary Fig.S3 and Fig.S4).

Transcriptome analysis of four different spatial compartments of the swarming model allowed to highlight the sequential gene regulations taking place during bacterial surface colonisation. A huge divergence in gene expression is observed between the MC and the three other compartments, including its adjacent compartment BS. These compartments govern metabolically active cells displaying a high upregulation of genes involved in several functions, essentially related to active growth: replication and division (*dnaAN* and *ftsEX* operons, *divIVA*), transcription and translation (*rpoA, pur* and *pyr* operons, *tRNA, rRNA*, ribosomal proteins genes), energy metabolism (glycolytic genes, thiamin and biotin biosynthesis), transport (several genes encoding transporters of various carbon and nitrogen sources such as amino acids, or transporters of different metal ions), motility and chemotaxis (*fla/che* operon, *hag, ycdA*). On the other flip, genes related to sporulation and gluconeogenic carbon metabolism are more expressed in the MC compared to the BS, DT and TP compartments, indicating that stress signals such as nutrient depletion initiated the sporulation process to face the harsh environmental conditions. Matrix related genes are more upregulated in the MC and the BS compared to the DT and TP, which is clearly observed by confocal imaging of the distribution of these subpopulations during a swarm (Supplementary Fig.S6). The regulations of these latter genes indicate that cells in the TP are more exploring the environment to incorporate nutrients from the medium, rather than expressing proteins involved in biofilm formation or sporulation, contrary to cells in the BS. Thus, each compartment is formed by cells under different physiological states with higher cellular heterogeneity going toward the MC.

Between adjacent static liquid compartments, half of the genome is differentially expressed with two biofilm populations coexisting in the same well. The DC (compartment between the two biofilms) have been long considered as a state similar to the planktonic population. Phenotypic and transcriptomic studies on various bacterial species, such as *Klebsiella pneumoniae* or *Streptococcus pneumoniae*, have shown that detached cells exhibit different gene expression patterns, distinct from both sessile and planktonic lifestyles^46,47,48,49^. Our results with *B. subtilis* confirm these previous observations. The transcriptome profile of the DC revealed a distinct state compared to both EX and ST planktonic phases as well as to the two biofilms in the static liquid model. For instance, the major matrix genes of the *tapA* and *epsA-O* operons, are downregulated in the DC compared to both the ST and the EX phases. These operons are more expressed in the DC and the PL than in the SB.

A spatio-temporal monitoring on the submerged level, revealed patterns of gene expression linked to the phenotypic heterogeneity observed during the different stages of biofilm development. We could highlight the heterogeneous expression of the different matrix genes (*epsA-O, tapA, bslA, srfAA, ypqP*, and *capB-E*) over both spatial and temporal levels. EPS and TasA are highly produced in the first few hours of incubation during the adhesion and development of the biofilm to the surface. The latter matrix components are assembled by BslA, required for biofilm architecture and biofilm hydrophobicity of the colony and pellicle^32,50^. In a previous study, we reported that *bslA* inactivation had an impact on the 3D structure of the colony and also on the stability of the pellicle, while no effect was observed for the submerged biofilm compartment after 24 hours of incubation^6^. 4D-CLSM allowed to demonstrate that *bslA* is expressed during the first 3 hours of SB development (together with the *epsA-O* and *tapA* operons) and then again in late stage of biofilm maturation after 17 hours, when the biofilm is already formed. A strong correlation between biofilm development and surfactin production was suggested within different *Bacillus* species. For instance, in *B. velezensis* FZB42, *B. amyloliquefaciens* UMAF6614 and *B. subtilis* 6051 defect in surfactin production has been shown to cause partial or severe biofilm defects^51,52,9^. On the other hand, in the *B. subtilis* strains NCIB3610 and NDmed, surfactin operon mutation was reported not to have any effect on biofilm formation (pellicle, colony and submerged)^6,53^. As an external signal, surfactin induces cells to express matrix genes^1^. The *srfAA* gene is expressed mainly in a temporal window between 21 and 36 hours during biofilm incubation after which one can re-observe expression of the EPS and TasA. In addition, the late expression of *ypqP* and *capE* observed in our imaging data is consistent with the small effect on biofilm formation at 24 hours previously reported for inactivation of these genes^6^. Only a subpopulation of the SB expresses the different matrix genes. Hence, heterogeneous spatio-temporal expression of matrix genes indicates specific requirements of the expensive matrix products through the different stages of biofilm development.

In a medium containing carbon and nitrogen excess, a major overflow pathway takes place through the conversion of pyruvate to acetate by the phosphotransacetylase-acetate kinase pathway to generate ATP. This pathway is positively regulated by a major regulator for the carbon metabolism CcpA which for instance activates the *ackA* gene, encoding an acetate kinase^54^. In this study we could see that *ackA* and the *cggR-gapA* operon (reporting glycolysis) were highly expressed during the first 13 hours, before being gradually downregulated, indicating that carbon source started to be limited afterwards (Fig. 2 and Fig. 4). Interestingly, this corresponded with the beginning of the first wave of dead cells that was clearly observed after 13 hours (Fig. 5), followed by the initiation of sporulation reported by *spoIIGA* (Fig. 2). Hence, these observations suggest that carbon source limitation triggered cell death which by turn provided carbon source for the initiation of the irreversible sporulation process. Cell death is also followed by competent and cannibal cell types, tagged by *comGA* and *skfA* genes, overexpressed at around 20 hours (Fig. 2), pointing out the capability of these cells to uptake exogenous DNA from the medium and produce spore-killing factors, allowing to delay their entry into the irreversible process of sporulation^19^. After the first wave of dead cells a slight increase in the live population was observed (Fig. 5, Supplementary Movie S1), which accommodates different subpopulations expressing either glycolysis or gluconeogenesis. Another expression of the *hag* motility gene is observed after 24 hours in a small subpopulation of the SB. This could correspond to pore forming swimmer cells as previously observed^44^. Surfactin, reported by *srfAA*, is overproduced around the same spatio-temporal window. Surfactin is involved in genetic competence and triggers matrix production^5,31,55,56^, in accordance with the upregulation of the genes *epsA-O, tapA, bslA, ypqP, capB-E* and *comGA* after 24 hours of incubation. Motility could also allow to increase the diffusion and activity of exoproteases (product of the *aprE* gene, among others) within the matrix biofilm. Moreover, cells undergoing sporulation are also present at that time as indicated by the overexpression of late sporulation genes (such as *spoVC*).

A highly structured colony has wrinkles, formed by mechanical forces due to increased cell density. Dead cells localised under these wrinkles, at the base of the biofilm and near the agar, lead to formation of channels that facilitate liquid transport within the biofilm^57,58^. In the SB, *B. subtilis* dead cells are clustered mainly on the top of the biofilm appearing in two waves during 13-24 hours and after 42 hours. The second wave of dead cells (after 42 hours) is also in the same accordance with the high expression of gluconeogenic genes (reported by *PgapB-gfpmut3*, Fig. 4). Despite that the DC at 24 hours are distinct from the planktonic populations (EX and ST) but are still in a physiological state closer to cells in EX rather than ST ones (Fig. 1b). This is illustrated by the extremely strong upregulation (around 9log2FC) of the *pstS-BB* and *tuaA-H* operons in the ST compared to the DC or the EX. These operons, involved in high-affinity phosphate uptake and teichuronic acid biosynthesis, respectively, are induced upon phosphate starvation^59^, which indicates that DC, like cells in the EX, do not suffer such conditions, contrary to cells in ST.

This report presents the first comparative description of the transcriptomic profiles of nine spatio-physiological populations of *B. subtilis* captured on solid, semi-solid and liquid cultures using the same strain and nutrient source. It allowed us to specify the singularities of each biofilm compartment and to pinpoint the fineness of their spatio-temporal regulation down to the single scale. The presented data give novel insights on the development and dispersal of *B. subtilis* surface-associated communities, with a special comprehension for the relation between central carbon metabolism regulation and dead cells on the submerged level (Fig.6). All the provided results summarised could serve as a unique resource for future studies on biofilm physiology to further investigate genetic determinants required for its control.

## METHODS

### Bacterial strains and growth conditions

The *B. subtilis* strains used during this study are listed in Table 1. NDmed derivatives were obtained by transformation with various plasmids or chromosomal DNA of various strains to introduce the corresponding suitable reporter fusion. The transcriptional fusions of the *gfpmut3* gene to the *ackA, hag, bslA, srfAA* or *gapB* promoter were constructed previously within the pBSB2 plasmid (pBaSysBioII) using ligation-independent cloning^60^, prior to integration into the chromosome of BSB168 in a non-mutagenic manner, resulting in strains BBA0093, BBA0231, BBA0290, BBA0428 and BBA9006, respectively; chromosomal DNA of each strain was used to transfer the corresponding fusion into NDmed by transformation. Similarly, fragments corresponding to the promoter regions of *epsA, ypqP, ctaA, narG, skfA, comGA, aprE, cggR, spoIIGA, spoVC*, and *tapA*, or to a region in the 3’ part of *capE*, were amplified by PCR from genomic DNA using appropriate pairs of primers (Supplementary Table S1). These fragments were inserted by ligation-independent cloning in pBSB2 or in pBSB8, a pBSB2 derivative with the *gfpmut3* and *spec* (spectinomycin resistance) genes replaced by *mCherry* (codon-optimised for *B. subtilis*) and *cm* (chloramphenicol resistance), respectively. The resulting plasmids were then used to integrate each corresponding transcriptional fusion into the chromosome of *B. subtilis* through single recombination. Transformation of *B. subtilis* was performed according to standard procedures and the transformants were selected on Luria-Bertani (LB, Sigma, France) plates supplemented with appropriate antibiotics at the following concentrations: spectinomycin, 100µg/mL; chloramphenicol, 5μg/mL. Before each experiment, cells were cultured on Tryptone Soya Agar (TSA, BioMérieux, France). Bacteria were then grown in synthetic B-medium composed of (all final concentrations) 15mM (NH_4_)_2_SO_4_, 8mM MgSO_4_.7H_2_O, 27mM KCl, 7mM sodium citrate.2H_2_O, 50mM Tris/HCl (pH 7.5), and 2mM CaCl_2_.2H_2_O, 1μM FeSO_4_.7H_2_O, 10μM MnSO_4_.4H_2_O, 0.6mM KH_2_PO_4_, 4.5mM glutamic acid (pH 8), 862μM lysine, 784μM tryptophan, 1mM threonine and 0.5% glucose were added before use^61^. Cultures for planktonic inoculum were prepared in 10mL B-medium inoculated with a single colony and shaken overnight at 37°C. The culture was then diluted to an OD_600nm_ of approximately 0.1 and grown at 37°C until it reached an OD_600nm_ of approximately 0.2. The procedure was repeated twice and finally the culture was grown to reach stationary phase, which was then used to inoculate swarming and liquid biofilm assays (Fig. 1).

### Swarming culturing condition

The OD_600_ was measured and the culture was diluted, and 2μL of diluted bacterial culture (adjusted to an OD_600nm_ of 0.01, ∼10^4^ CFU) were inoculated at the centre of B-medium agar plate and incubated for 24 hours at 30°C with 50% relative humidity. Plates (9cm diameter, Greiner bio-one, Austria) containing 25mL agar medium (0.7% agar) were prepared 1 hour before inoculation and dried with lids open for 5 minutes before inoculation.

### Liquid biofilm culturing condition

Cultures were performed in microplates, either 3mL in 12-well microplate (Greiner bio-one, Germany) or 150µL in 96-well microscopic grade microplate (µclear, Greiner bio-one, Germany), inoculated from a stationary phase culture and adjusted to an OD_600nm_ of 0.01. The plates were incubated at 30°C for 24 hours, followed by either local cell harvesting or microscopic imaging. The 96-well plate was used for kinetic monitoring of the submerged biofilm, and the pellicle was collected from a 12-well plate for observations. When necessary, the medium was supplemented with 200μM isopropyl-β-d-thiogalactopyranoside (IPTG) to induce Gfp expression from the *Phyperspank* promoter.

### Local mesoscopic cell harvest for RNA-seq

For EX and ST phases (OD_600_ ∼0.6 and ∼2.8, respectively), 6mL of each culture were collected and pelleted by centrifugation at 8,000 × g at 4°C for 30 seconds. The pellet was then homogenised by 500µl TRIzol reagent (Invitrogen, Carlsbad, CA, USA) for stabilising the RNA in the cell. For the swarming model, using 24hr plates, four spatially localised compartments (MC, BS, DT and TP) were collected independently. Collection was done manually by using a scraper (SARSTEDT, USA) starting from the tips down to reach the mother colony (that was collected by a loop). Cells of each localised compartment were collected from 16 plates in an Eppendorf tube (CLEARline microtubes, Italy) containing 500µL TRIzol reagent. For a 24hr static liquid model, 6 wells (from a 12-well microplate) were used to collect each sample. By using a scraper, PL were collected in 6mL water. For DC, 1ml from the supernatant was collected from 6 wells. For the SB collection, after discarding all the rest of the liquid, 1ml water was added in a well and cells were collected by scratching with a pipet tip. Samples were centrifuged rapidly for 30 seconds (8,000 × g at 4°C) and pellets resuspended in 500µL Trizol.

A centrifugation step for all the above collections for 1 minute to discard the TRIzol reagent was done and samples were snap-frozen by liquid nitrogen to be transferred to −80°C ready for the RNA extraction step. For each of the 9 samples, 3 biological replicates were done.

### RNA extraction for RNA-seq

For all nine different conditions, a washing step for the pellets of the *B. subtilis* NDmed was done with 1mL TE (10mM Tris, 1mM EDTA, pH=8) + 60µl 1 M EDTA followed by centrifugation for 30 seconds (8,000 × *g* at 4°C). Cell pellets were suspended in 1mL TRIzol reagent. Cell suspension was transferred to a Fastprep screw cap tube containing 0.4g of glass beads (0.1mm). Cells were disrupted by bead beating for 40 seconds at 6.5m/s in a FastPrep-24 instrument (MP Biomedicals, United states). The supernatant was transferred to an Eppendorf tube and chloroform (Sigma-Aldrich, France) was added in a ratio of 1:5, followed by centrifugation at 8,000 × *g* for 15 minutes at 4°C. The chloroform step was repeated twice. The aqueous phase was transferred to new Eppendorf, where sodium acetate (pH=5.8) was added to a final concentration of 0.3M and 500µl of isopropanol (Sigma-Aldrich, France). Samples were left overnight at −20°C and then centrifuged for 20 minutes. Pellets were washed twice by 75% of ethanol (VWR, France) followed by centrifugation for 15 minutes at 4°C. Then pellets were dried for 5 minutes under the hood. A RNA cleanup kit (Monarch RNA Cleanup Kit T2050, New England Biolabs, France) was used to further clean the RNA samples. Extracted RNA samples were stored in water RNAase/DNAse free (Ambion, United Kingdom) at −80°C. Nanodrop and Bioanalyzer instruments were used for quantity and quality controls. Library preparation including ribosomal RNA depletion and sequencing was performed by the I2BC platform (Gif-sur-Yvette, France) using TruSeq Total RNA Stranded and Ribo-Zero Bacteria Illumina kits, an Illumina NextSeq 550 system and NextSeq 500/550 High Output Kit v2 to generate stranded single end reads (1 × 75bp).

### RNA-seq data analysis

Primary data processing was performed by I2BC platform and consisted of: demultiplexing (with bcl2fastq2-2.18.12), adapter trimming (Cutadapt 1.15), quality control (FastQC v0.11.5), mapping (BWA v0.6.2-r126, ^64^) against NDmed genome sequence (NCBI WGS project accession JPVW01000000, ^16^). This generated between 13M and 29M of uniquely mapped reads per sample which were summarised as read counts for 4028 genes (featureCounts, ^65^) after discarding 7 loci whose sequences also matched External RNA Controls Consortium (ERCC) references. The downstream analysis was performed using R programming language. Samples were compared by computing pairwise Spearman correlation coefficients (ρ) and distance (1-ρ) on raw read counts which were summarised by a hierarchical clustering tree (average-link). Detection of DEGs used R package “DESeq2” (v1.30.1, ^66^) to estimate p-values and log2 fold-changes. To control the false discovery rate, for each pair of conditions compared, the vector of p-values served to estimate q-values with R package “fdrtool” (v1.2.16, ^67^). DEGs reported for pairwise comparisons of *B. s*ubtilis spatial compartments were based on a q-value≤0.05 and, unless stated otherwise, |log2FC|≥1. Fragment counts normalised per kilobase of feature length per million mapped fragments (fpkm) computed by DESeq2 based on robust estimation of library size were used as values of expression levels for each gene in each sample. Genes were compared for their expression profiles across samples for selected sets of conditions based on pairwise Pearson correlation coefficients (r) and distance (1-r) computed on log2(fpkm+5) and average-link hierarchical clustering of the distance matrix. Accordingly, the associated heatmaps represent gene-centred variations of log2(fpkm+5) values across samples. Gene clusters defined by cutting the hierarchical clustering trees at height 0.3 (corresponding to average r within group of 0.7) were numbered by decreasing number of the genes coupled in the same group, G1 for the largest. The resulting gene clusters were systematically compared to *Subtiwiki* functional categories and regulons^38^ (from hierarchical level 1 to level 5) using exact Fisher test applied to 2×2 matrices. The results of the comparisons with *Subtiwiki* functional categories were summarised in the form of stacked bar plots after manually assigning each gene to the most relevant category in the context of this study (when the same gene belonged to several categories) and a grouping of categories corresponding to hierarchical level 2 excepted for “Metabolism” (level 1), and “motility and chemotaxis” and “biofilm formation” (level 3). The whole transcriptomic data set has been deposited in GEO (accession number <u>GSE214964</u>).

### CLSM

The biofilm models were observed using a Leica SP8 AOBS inverted laser scanning microscope (CLSM, LEICA Microsystems, Wetzlar, Germany) at the INRAE MIMA2 platform (https://doi.org/10.15454/1.5572348210007727E12). For observation, strains were tagged fluorescently in green with SYTO 9 (0.5:1000 dilution in water from a stock solution at 5µM in DMSO; Invitrogen, France) and SYTO 61 (1:1000 dilution in water from a stock solution at 5µM in DMSO; Invitrogen, France), a nucleic acid marker. After 15 to 20 minutes of incubation in the dark at 30 °C to enable fluorescent labelling of the bacteria, plates were then mounted on the motorised stage of the confocal microscope. For the carbon metabolism reporting genes, the 3D (xyz) acquisitions were performed by a HC PL FLUOTAR 10x /0.3 DRY objective (512 × 512 pixels, pixel size 0.361 µm, 1 image every z = 20 µm with a scan speed of 600 Hz, and a pinhole70µm) to be able to capture the submerged and the pellicle in the same well. Moreover, the different selected compartments were scanned using either HC PL APO CS2 63x/1.2 water immersion or 10x objective lenses. SYTO 9, Gfp and IP excitation was performed at 488 nm with an argon laser, and the emitted fluorescence was recorded within the ranges 500-550 nm and 600–750 nm, respectively on hybrid detectors. SYTO 61 or mCherry excitation was performed at 561 nm with an argon laser, and the emitted fluorescence was recorded within the range 600–750 nm on hybrid detectors. The 3D (xyz) acquisitions were performed (512 × 512 pixels, pixel size 0.361 µm, 1 image every z = 1 µm with a scan speed of 600 Hz). For 4D (xyzt) acquisitions an image was taken every 1 hour for 48 hours or 1 and half hours for 72 h.

The whole 4D-CLM data set has been deposited in Recherche Data Gouv (https://doi.org/10.57745/Z511A6).

### Image analysis

Projections of the biofilm, 3D or 4D were constructed from Z-series images using IMARIS 9.3 (Bitplane, Switzerland). Space-time kymographs were constructed with the BiofilmQ visualisation toolbox from 4D-CLSM series^68^.

## ACKNOWLEDGEMENTS

This work was supported by INRAE. Y. Dergham is the recipient of fundings from the Union of Southern Suburbs Municipalities of Beirut, INRAE, Campus France PHC CEDRE 42280PF and Fondation AgroParisTech. P. Sanchez-Vizuete was the recipient of a PhD grant from the Région Ile-de-France (DIM ASTREA). Pr. M. Jules (University Paris-Saclay, AgroParisTech, INRAE) is acknowledged for the gift of plasmids pBSB2 and pBSB8, and of strains BBA0093, BBA0231, BBA0290, BBA0428 and BBA9006. M. Calabre (INRAE) is acknowledged for technical assistance. We acknowledge the sequencing and bioinformatics expertise of the I2BC High-throughput sequencing facility, supported by France Génomique (funded by the French National Program “Investissement d’Avenir” ANR-10-INBS-09). Biofilm imaging was realized at the INRAE MIMA2 imaging platform https://doi.org/10.15454/1.5572348210007727E12. This work is performed under the umbrella of the European Space Agency Topical Team: Biofilms from an interdisciplinary perspective.

## Author contributions

Y.D., D.L.C, K.H. and R.B. designed research; Y.D., D.L.C, E.H., J.D. and P.S.V performed research; Y.D., P.N., D.L.C and R.B analysed data; Y.D., D.L.C, P.N., K.H. and R.B. wrote the manuscript with support from all authors.

