## Supplementary material for "Multi-scale transcriptome unveils spatial organisation and temporal dynamics of *Bacillus subtilis* biofilms"

**1- Supplementary Figures**

**2- Supplementary Movies**

**3- Supplementary Table**

### 1- Supplementary Figures

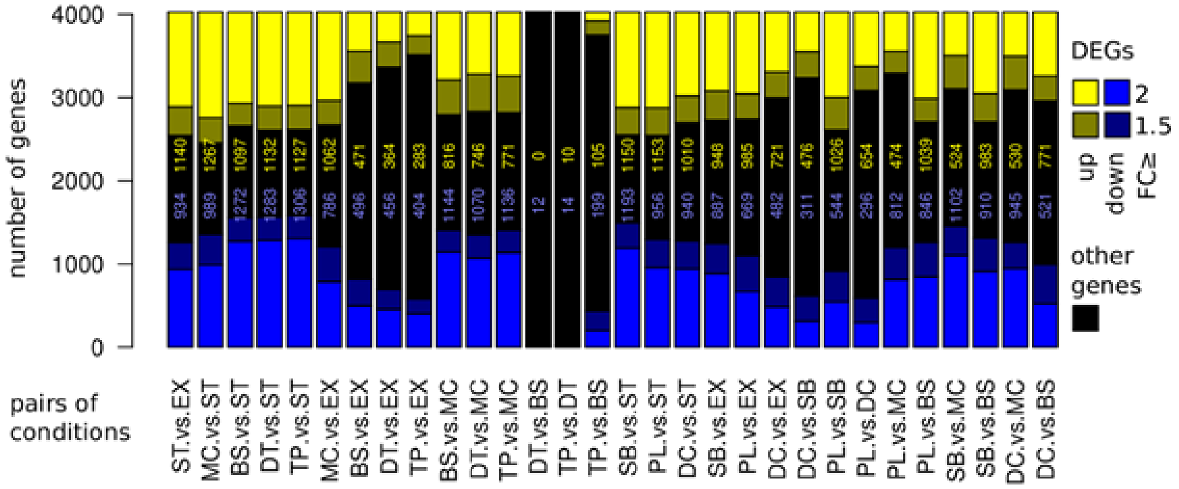


***Figure S1: Differentially expressed genes (DEGs) comparisons across the different compartments of B. subtilis in the RNAseq analysis.*** *Out of 4028 genes, the numbers of DEGs in pairwise comparison of different B. subtilis spatial compartments (q-value≤0.05, | log2FC≥1) are reported in yellow (up-regulated) and in blue (down-regulated). For instance, comparison of the stationary to the exponential phase (ST vs. EX) shows that 1140 and 934 genes are significantly upregulated and downregulated with a minimum log2FC, respectively. Among the 1140 upregulated genes, around 36% are related to sporulation, biofilm formation (tasA and epsA-O operons, as well as bslA), coping with stress, or carbon metabolism, and 30% to other minor functions; the remaining (approximately 34%) are of unknown function, e.g. ydhF, ywcI, yycOPQ, and yjdB, which show the highest upregulation, all above 7log2FC. Comparison between adjacent compartments of a swarm highlights some genes that are differentially expressed. Between the base and the mother colony (BS vs MC) there are 815 genes upregulated and 1140 genes downregulated. In the dendrites, 12 genes are downregulated compared to the base (DT vs BS), ppsD-E, rpsNB, yxeG, yrpE, and the folEB-yciB, znuACB and pftAB operons. In the tips when compared to the dendrites (TP vs. DT), 14 genes are downregulated (veg, ykoY, ydzJ, spoVS, yydF, sspF, yqzM, tRNA-Met, tRNA-Thr, tRNA- Ser, tRNA-Asp, tRNA-Val, and 2 genes tRNA-Arg) and 10 are upregulated (tRNA-His, rbsD, melRE, ldh, cydA, and the ydgGH and ytbDE operons). The ycdA gene, encoding a lipoprotein required for swarming motility, shows gradual upregulation by a log2FC from one localised compartment to its adjacent one going from the mother colony to the tips. Even though between very close compartments only a few genes display some differential changes, 105 genes are upregulated in the tips compared to the base (TP vs. BS) and 199 genes are downregulated. For the liquid culture, 654 genes are upregulated and 296 are downregulated in the pellicle compared to the detached cells (PL vs. DC). Moreover, 476 genes are upregulated and 311 genes downregulated in the detached cells compared to the submerged (DC vs SB).*

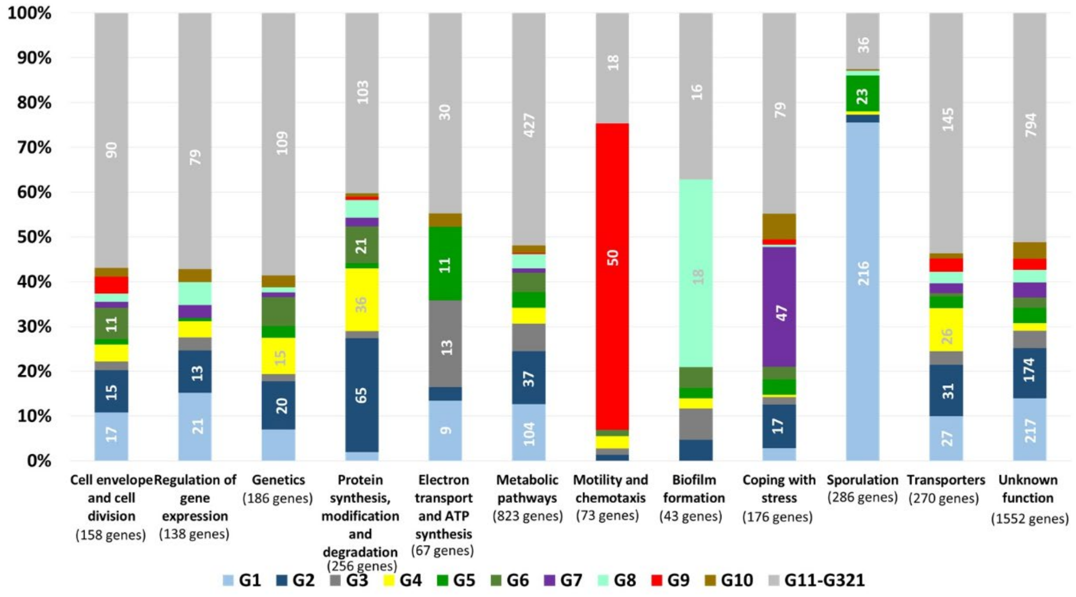


***Figure S2: Relative distribution for the expression clusters in Figure 1c for 4028 genes according to the Subtiwiki-derived functional categories.*** *The groups are clustered depending on the similarity of their gene expression profiles and named depending on the number of genes coupled in the same group by decreasing order. Group 1 (G1) governs the highest number of genes, 634 genes, differentiated along the different conditions, from which 216 genes are required for sporulation. Biofilm formation related genes, eps and the tapA operons with the slrR gene, are governed in group 8 (G8), while matrix and chemotaxis related genes, mainly the fla/che operon, is governed in group 9 (G9). Around 1552 out of 4028 (~38%) of the genes are either unknown or poorly characterised.*

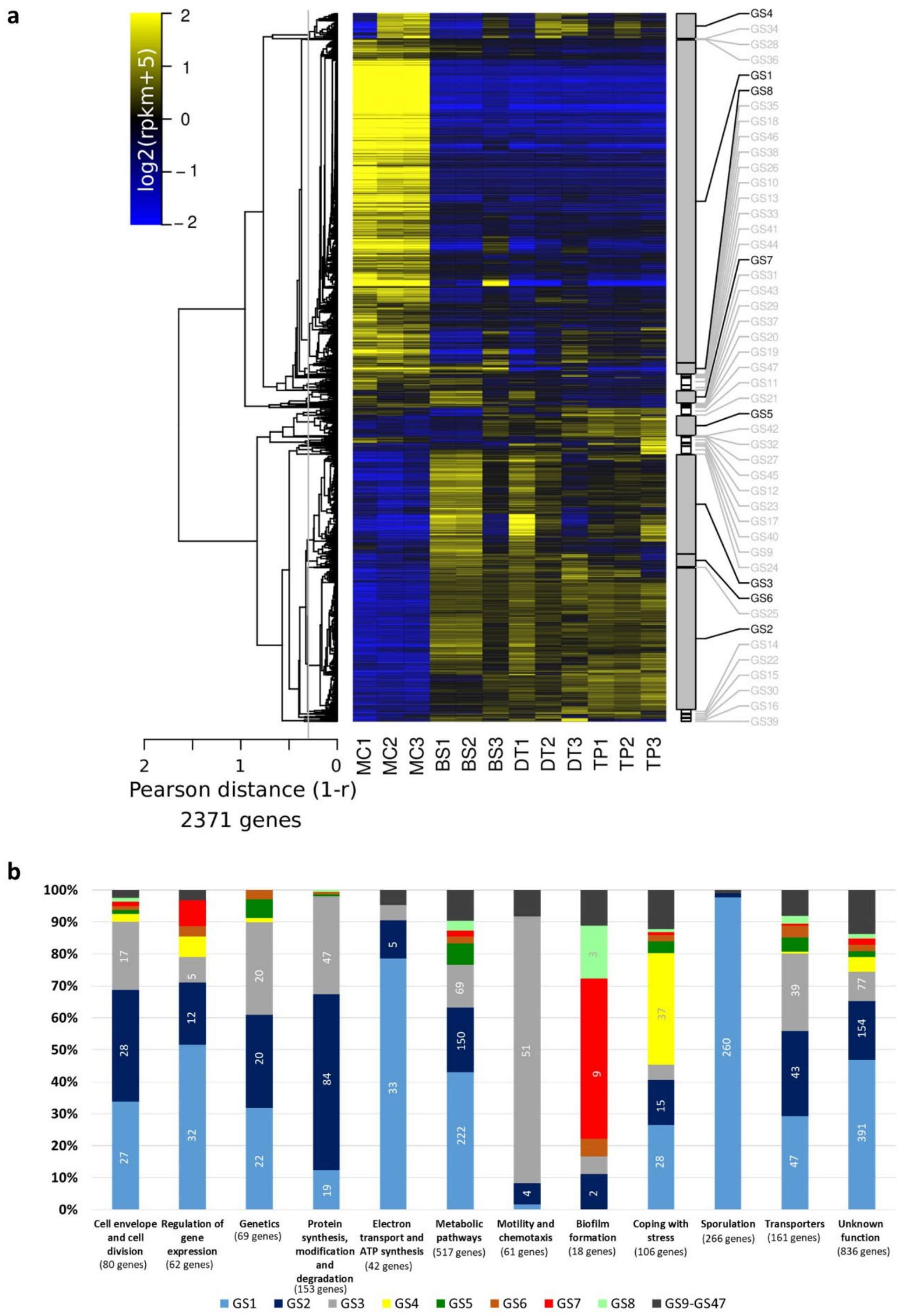


***Figure S3: Transcriptome remodelling of differentially expressed genes during swarming.*** *(a) Heatmap representation of the relative variations of expression level across samples for DEGs (2371 out of the 4028) identified in the 6 pairwise comparisons of 4 different localizations (MC, BS, DT, and TP). The colour code reflects the comparison to the mean computed for each gene across the 12 samples (log2 ratio). The hierarchical clustering tree shown on the left side of the heatmap (average link) was cut at average Pearson correlation of 0.7 (vertical grey line) to define the expression clusters shown on the right side of the heatmap. Clusters were named (from GS1 to GS47) by decreasing sizes and those containing more than 30 genes are highlighted (name printed in black). (b) Distribution into expression clusters for genes in Subtiwiki-derived functional categories.*

**Additional interpretation for the RNAseq data for Figure S3:**

**A retrospective view on the adjacent spatial compartments of a swarm shows differential gene expression**

A careful examination of global transcription in the swarming model would allow us to better understand the sequential gene regulations occurring through surface colonisation to biofilm formation. From the 4028 genes of the *B. subtilis* NDmed, 2371 genes are differentially expressed between the four localised compartments of a swarm and grouped according to the similarity of their gene expression, represented as a heatmap in Figure S3a. There are 47 groups of genes differentially expressed within the swarming model (GS1 to GS47). Figure S3b, represents a functional category percentage of the 2371 genes across the different conditions of a swarm.

GS1 is a large group of genes (1082) highly expressed in the mother colony and repressed through the swarm (Fig. S3a), corresponding to the different functions occurring at the same time in this biofilm model. Approximately 80% of the known genes are related to the functional group of the 'Electron transport and ATP synthesis' from which 22 out of 33 are genes encoding functions related to respiration *i.e. fnr*, *arfM, narG-I, cydBCD, nasD, cccA, qcrABC, ccdA, ctaC-G,* and *ythAB* with a significant upregulation of minimum Log2FC in the mother colony compared to the base. Moreover, 222 genes encode other metabolic pathways related to carbon metabolism (*i.e. yjmCDF* with 8Log2FC upregulated in the MC compared to the BS, *acoAB* with a 7Log2FC, *licABCH* with a 4Log2FC, *gapB* with a 4Log2FC, *pckA* with a 3Log2FC, *alsDS* with a 2Log2FC), lipid metabolism (*fadA-R* with a 4Log2FC), nucleotide metabolism (*pucH* with a 3Log2FC), miscellaneous metabolic pathways (*glgA* with a 5Log2FC and *ntdABC* with a 7Log2FC), and nitrogen metabolism (*mmgA-F* with approximately 6Log2FC), and genes related to coping with stress (e.g. 5Log2FC for *skfA* and a 6Log2FC for *oxdC*). Sporulation is the function that is majorly differentially expressed in this group by 98%, for example, 5Log2FC for both *spoIIGA* and *spoVD*, 2Log2FC for *ypqP*, upregulated in the mother colony compared to the base.

The heatmap (Fig. S3a) shows an upregulation in both the mother colony and the base in both swarming groups, GS7 and GS8. These groups contain many genes involved in biofilm formation (Fig. S3b). Genes of the *eps* operon (*epsF-N)* and *slrR* are in GS7, the *tapA* operon (*tapA-sipW-tasA*) is in GS8, and all are significantly downregulated by Log2FC in the tips compared to the base (TP vs. BS). A comparison between the tips and the mother colony (TP vs MC) shows a slight downregulation of the *eps* and *slrR* genes (less than a Log2FC), however, the *tasA* operon is downregulated by approximately 1.5Log2FC. No significant difference in gene expression is observed between the adjacent compartments of a swarm (through the different comparisons BS vs. MS, DT vs. BS, or TP vs DT). Genes related to motility and chemotaxis present in GS3 and GS2 (representing 90% of this functional category) are downregulated in the mother colony and upregulated during the swarm. The *fla/che* operon is upregulated by 2Log2FC, in which the *hag* gene is the highest differentially expressed by 5Log2FC in the tips compared to the mother colony (Tp vs. MC). Several genes poorly characterised or of unknown function like *ywdK*, *yhoC*, *yuzM*, *yodT*, *yqfX*, or *ydgB* are highly expressed (more than 7Log2FC) in the mother colony compared to the swarm, while on the other hand, *ylxF*, *yscB*, and *yxkC* are expressed with a 3Log2FC in the swarming compartments compared to the mother colony. An upregulation by 4Log2FC is observed for the *ydgGH* (probable) operon in the tips compared to the other swarming compartments.

Comparison between adjacent compartments of a swarm highlights some genes that are differentially expressed (Fig. 1b). Between the base and the mother colony (BS vs MC) there are 815 genes upregulated and 1140 genes downregulated, most of which have been described in the GS1. In the dendrites, 12 genes are downregulated compared to the base (DT vs BS), *ppsD-E*, *rpsNB*, *yxeG*, *yrpE*, *folEB-yciB* and the *znuACB* and *pftAB* operons. In addition, 14 genes are downregulated (*veg, ykoY, ydzJ, spoVS, yydF, sspF, yqzM, tRNA-Met, tRNA-Thr, tRNA- Ser, tRNA-Asp, tRNA-Val, and 2 genes tRNA-Arg*) and 10 upregulated (*tRNA-His, ytbE, ytbD, rbsD, melE, melR, ldh, cydA, ydgH and ydgG*) in the tips when compared to the dendrites (TP vs. DT). The *ycdA* gene, encoding a lipoprotein required for swarming motility, shows gradual upregulation by a Log2FC from one localised compartment to its adjacent one going from the mother colony to the tips.

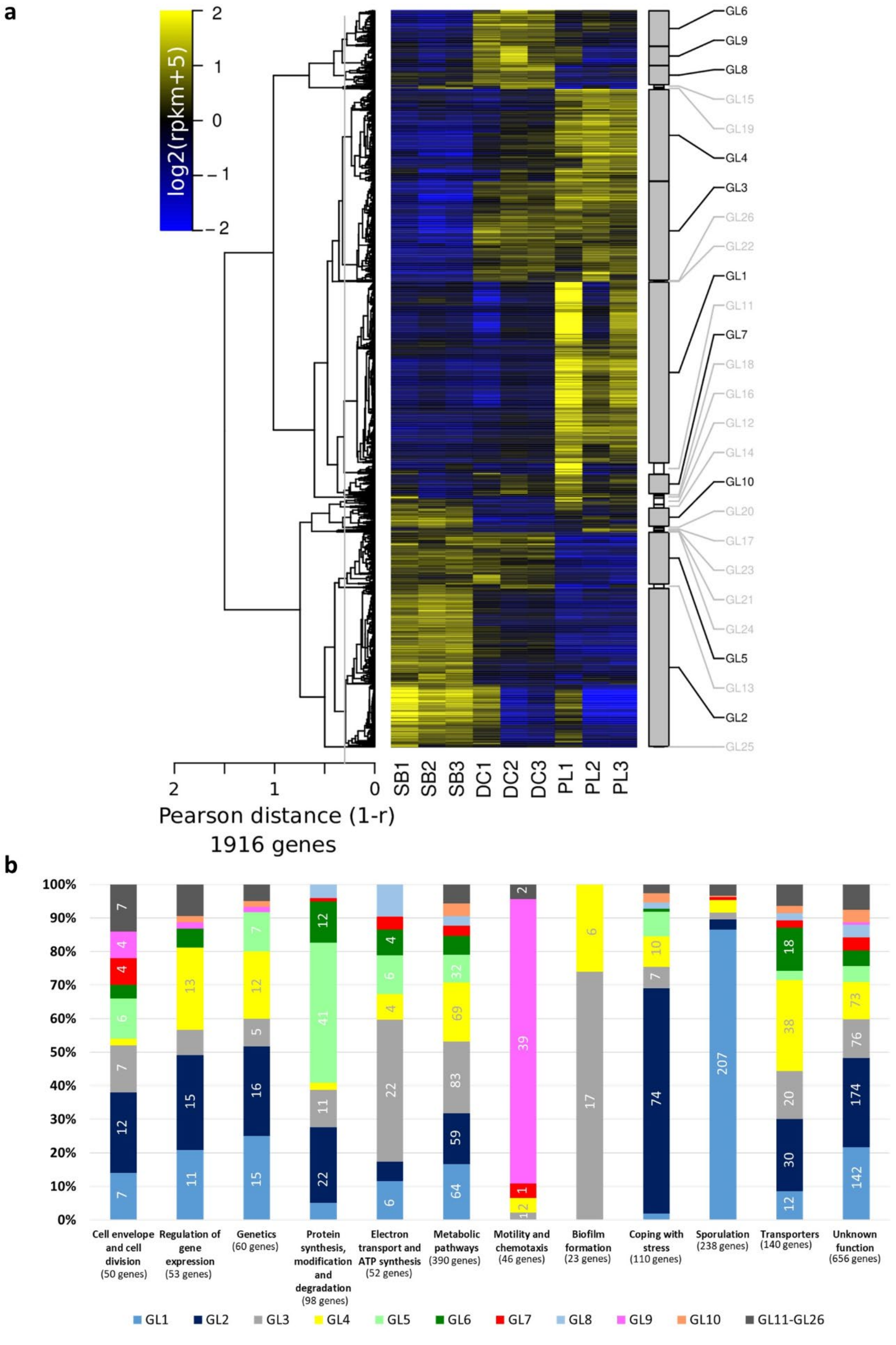


***Figure S4: Transcriptome remodelling shows that half of the genome is differentially expressed between the floating pellicle and the submerged biofilms coexisting in the same microplate well.*** *(a) Heatmap representation of the relative variations of expression level across samples for DEGs (1916) identified in the 3 pairwise comparisons of 3 different localizations (SB, DC, PL). Clusters were named (from GL1 to GL26) by decreasing sizes and those containing more than 30 genes are highlighted. (b) Distribution into expression clusters for genes in Subtiwiki-derived functional categories.*

#### Additional interpretation for the RNAseq data for Figure S4:

**Half of the genome is differentially expressed between the floating pellicle and the submerged biofilms coexisting in the same microplate well**

In this study, we have separately collected the submerged (SB), the pellicle (PL) as well as the detached cells (DC), a compartment between the SB and PL. From the 4028 genes, 1916 are differentially expressed between the three localised compartments clustered by 26 groups (GL1 to GL26) with functional category percentage, represented in Figure S4.

Groups GL6, GL9 and GL8 (Fig. S4a) shows an upregulation in the gene expression profile for the detached cells compared to the pellicle and the submerged. Classification by functional categories (Fig. S4b) shows that 85 % of the motility and chemotaxis genes are in group GL9. This group contains genes of the *fla/che* operon and *hag,* the latter showing an upregulation by approximately 3Log2FC and 1.5Log2FC in the detached cells compared to the submerged and the pellicle, respectively. Interestingly, belonging to the *fla/che* operon in GL9, *swrD* encodes a swarming protein (SwrD) used to promote flagellar power. The *swrD* gene is the highest down regulated gene in the pellicle compared to the detached cells by a 2Log2FC (PL vs. DC), which means it is a gene highly upregulated in the detached cells compared to the pellicle. This could suggest that the liquid culture is rather viscous, requiring a swarming-like process to efficiently migrate in the liquid column.

GL3 and GL4 groups contain most of the already known genes involved in biofilm formation (Fig. S4b), which are upregulated mainly in the pellicle (PL) and partially in the detached cells (DC) (Fig. S4a). Biofilm genes in these groups are the *tapA* and *epsA-O* operons, *dhbACB*, *slrR*, *sinI*, *bslA*, and *spo0A*. Comparison between the matrix genes for the two biofilm populations after 24 hours for the liquid model indicates that the *tapA* operon is upregulated by 4Log2FC, the *epsA-O* operon by approximately 3Log2FC, and *bslA* by a Log2FC in the pellicle compared to the submerged (PL vs SB). As for the regulators encoding genes in this group, *slrR* and *sinI* show an upregulation by 2LogFC and *spo0A* by a Log2FC in the pellicle compared to the submerged. Sporulation genes, of which about 87% are present in GL1 (Fig. S4), show a higher upregulation in the pellicle compared to the other two populations present, detached and submerged, like the *cge*, *cot*, *cwl*, *spoII*, *spoIII* , *spoIV*, *spoV* operons and genes. The most highly expressed genes in the pellicle compared to the submerged (PL vs. SB) by around 3Log2FC, are *ysxE*, *spoVID*, *spoIVA*, as those compared to the detached cells (PL vs. DC) are *cotC*, *yxeD*, *cotU*. The *ypqP* gene, potentially involved in the synthesis of polysaccharide, has been suggested to be involved in the addition of polysaccharides to the spore envelope. This gene is upregulated by 1.6Log2FC and 1.9Log2FC in the pellicle compared to the submerged or the detached cells, respectively.

GL5, a group clustering 135 genes, is upregulated in the detached cells (DC) and submerged (SB) (Fig. S4a). The *narG-I* operon, involved in nitrate respiration, is highly upregulated by a 5Log2FC in both detached and submerged cells compared to pellicle (PL vs. DC, and PL vs. SB). Carbon metabolism related genes, *i.e.*, *lctP*, *gapA*, *eno*, *cggR*, *ackA*, *pgm*… are all upregulated by more than Log2FC in the submerged compartments (SB and DC) compared to the pellicle (PL). Figure S4a, shows a high expression in the submerged population compared to the pellicle and the detached one in the GL2 clustering group. This group contains around 70% of genes related to the functional category coping with stress (Fig. S4b), most of which are regulated by the SigB regulon *i.e.*, *yjgD*, *ygxB*, *csbx*, *ydbD*, *ktaE*, *yhxD*,... that are upregulated by 3Log2FC and 2Log2FC in the submerged compared to the pellicle or to the detached cells, respectively (downregulated in the comparisons PL vs. SB and DC vs. SB).

About 34 % of the differentially expressed genes between the different populations of the liquid model are poorly characterised or of unknown function (Fig. S4b). Thus, in the comparison between the pellicle and the submerged (PL vs. SB), among others genes like *yoaC*, *yitJ*, and *ywcL* are upregulated while *yjgC*, *ydaC*, and *yxlE* are downregulated by 3Log2FC. Moreover, genes such as *ydjJ*, *yoxB*, and *yycD* are downregulated by more than 2Log2FC in the detached cells compared to the submerged (DC vs. SB). The *ywmC* and *ypzD* genes are upregulated by approximately 3Log2FC and *yclD*, *ybfA*, *yhbD*, and others are downregulated by more than 2Log2FC in the pellicle compared to the detached cells (PL vs. DC).

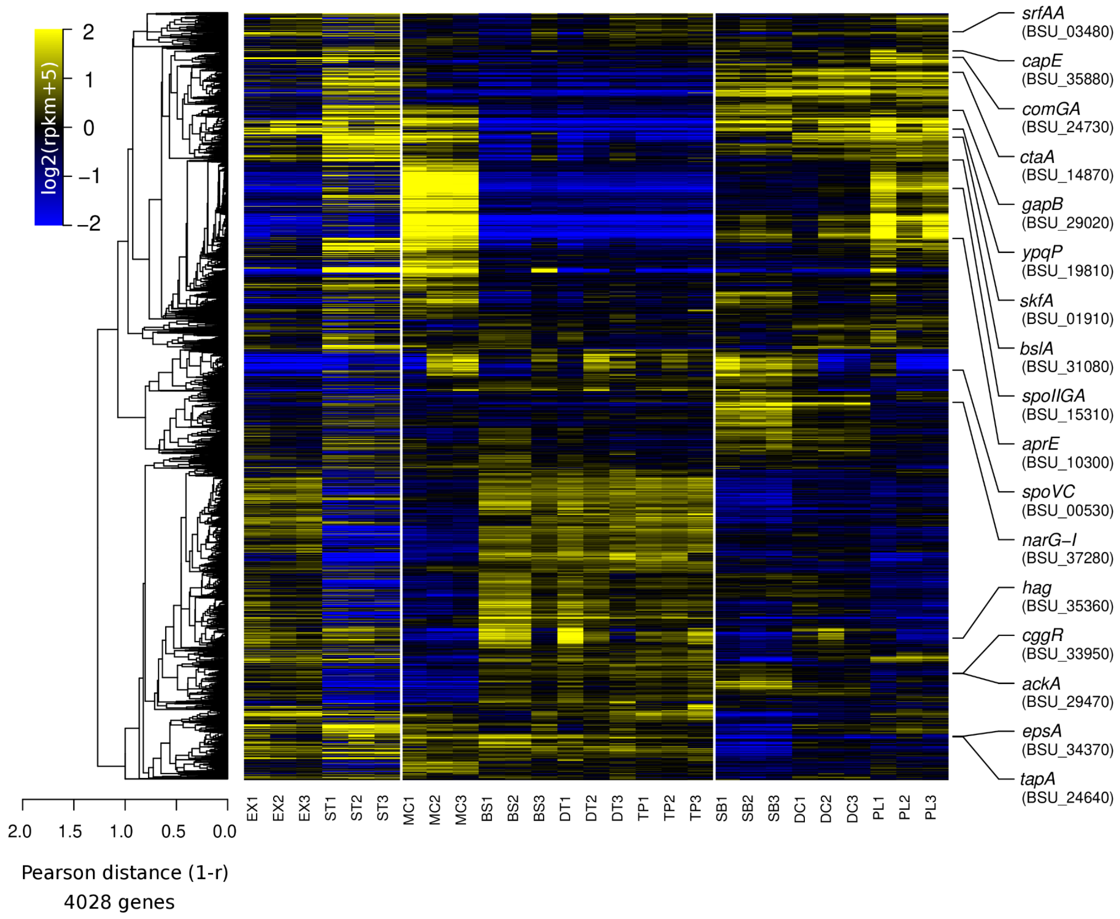


***Figure S5: Global heatmap for all the different conditions pinpointing reported genes that have been selected.*** *Global heatmap representation for the 4028 genes present in NDmed across the spatially selected surface-associated compartments. The colour code reflects the comparison to the mean computed for each gene (log2 ratio) taking as a reference the average of all conditions, except the planktonic ones (EX and ST). On the extreme right are the selected and transcriptionally reported genes, representatives of the different physiological activities potentially present in a biofilm.*

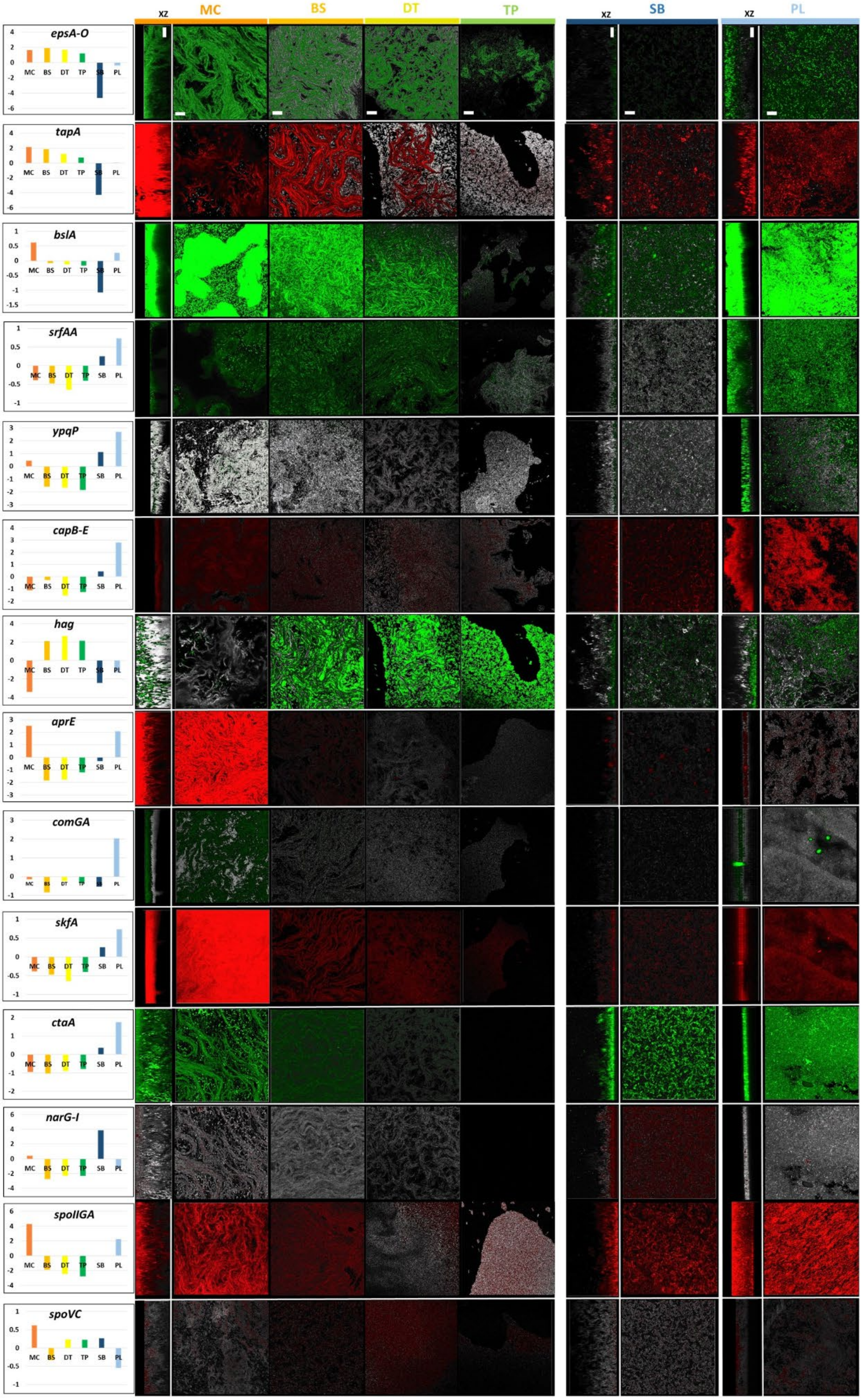
xz
MC
BS
DT
TP
xz
SB
xz
PL
epsA-O
tapA
bslA
srfAA
ypqP
capB-E
hag
aprE
comGA
skfA
ctaA
narG-I
spoIIGA
spoVC

***Figure S6: From mesoscopic to microscopic scale.*** *The RNAseq data is represented on the most left panels with a log2FC scale of an average of the three biological replicates compared to the average of all presented conditions together. The transcriptome results of epsA, capA and narG are used as representatives for their corresponding operon epsA-O, capB-E and narG-I, respectively. For confocal imaging, contrast was obtained by chemical staining or through expression of a second reporter gene fusion. For swarming plates (MC, Mother Colony; BS, Base; DT, Dendrites; TP, Tips) and static liquid cultures (SB, Submerged; PL, Pellicle), images represent a section (x and y 50µm and z 30µm); the scale bar represents 20µm. For the biofilm models (MC, SB and PL), a projection through the xz plane is presented on the left of each image; the vertical scale bars indicate the bottom level of the surface associated communities, in contact with the agar surface, solid surface or liquid surface, respectively. Images are representatives of the major phenotype from at least three replicates for each condition.*

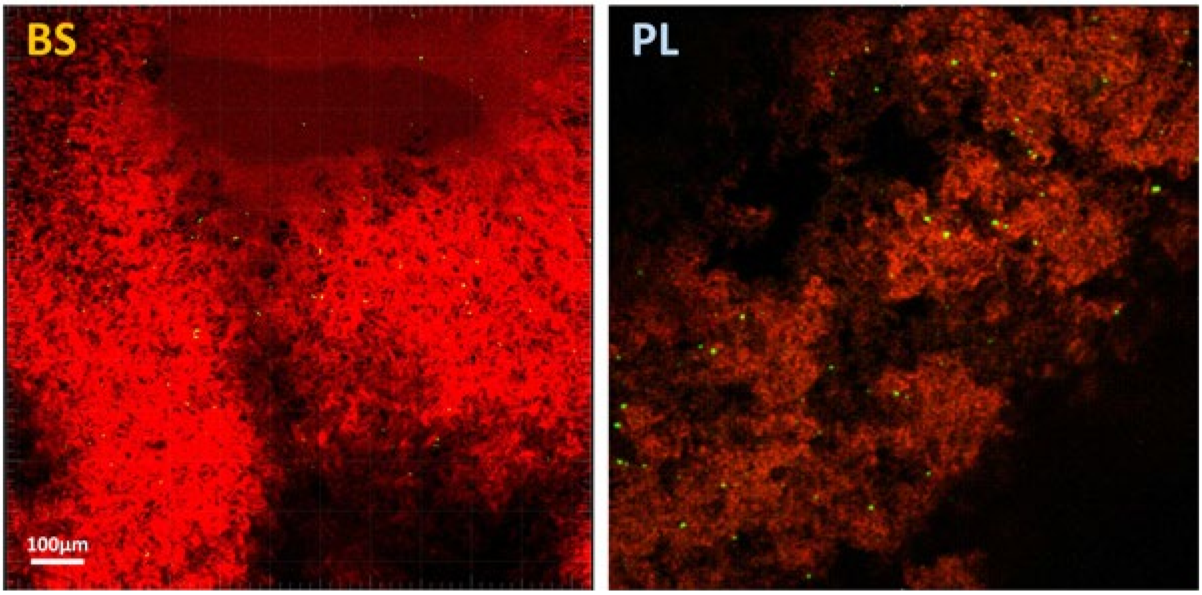


***Figure S7: In situ spatial monitoring unveils the switch from glycolytic to gluconeogenic regime at the cell level.*** *Spatial confocal imaging using strain GM3900 reporting transcription of cggR-gapA by mCherry (in red) and of gapB by Gfp (in green), for the BS and PL, from the swarming and static liquid models respectively, after 24 hours of incubation at 30°C. Same protocol as for the transcriptome analysis was used, except for the static liquid model the usage of 96-well microplate instead of the 12-well. Three replicative observations were performed independently for each model.*

#### 2- Supplementary Movies

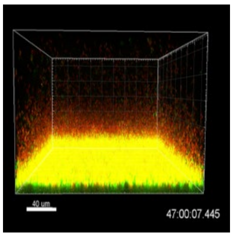


**Movie S1:** Submerged biofilm dynamics of *B. subtilis* NDmed-GFP (GM3649) with propidium iodide. The movie is representative of 3 replicates, with an image taken every 1 hour for 48 hours at 30°C, and represented by Imaris 4D projection.

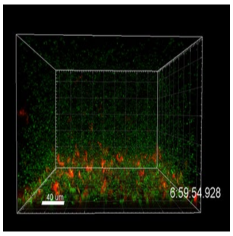


**Movie S2:** Spatio-temporal observation of strain GM3924, reporting transcription of *hag* by Gfp (in green) and of *tapA* by mCherry (in red) during the submerged biofilm development. (1 image every 1 hour for 48 hours, Imaris 4D projection representation).

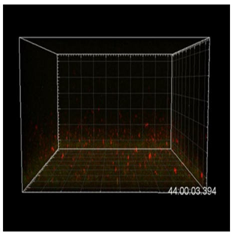


**Movie S3:** Spatio-temporal observation of strain GM3903, reporting transcription of *ackA* by Gfp (in green) and of *aprE* by mCherry (in red) during the submerged biofilm development. (1 image every 1 hour for 48 hours, Imaris 4D projection representation).

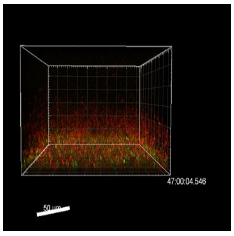


**Movie S4:** Spatio-temporal observation of strain GM3912, reporting transcription of *comGA* by Gfp (in green) and of *skfA* by mCherry (in red) during the submerged biofilm development. (1 image every 1 hour for 48 hours, Imaris Easy 4D projection representation).

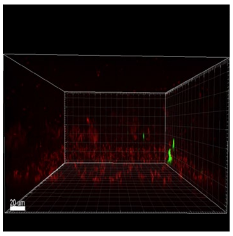

**Movie S5:** Spatio-temporal observation of strain GM3900, reporting transcription of *gapB* by Gfp (in green) and of *cggR* by mCherry (in red) during the submerged biofilm development. (1 image every 90 minutes for 72 hours, Imaris 4D projection representation).

#### 3- Supplementary Table

**TABLE S1:** Forward (F) and reverse (R) primers used for the construction of plasmids with reporter fusions.

| Fusion | Primers | Plasmid |
|---|---|---|
| *PepsA-gfpmut3* | F: CCGCGGGCTTTCCCAGCCCTTTAACCGATCATC<br>R: GTTCCTCCTTCCCACCTTCAGCCTTCCCGCG | pBSB2epsA |
| *PypqP-gfpmut3* | F: CCGCGGGCTTTCCCAGCTTGCCAAACTCATAAGAATG<br>R: GTTCCTCCTTCCCACCTCCAACTCTCGTTTCTCTAC | pBSB2ypqP |
| *PctaA-gfpmut3* | F: CCGCGGGCTTTCCCAGCGTAAGAAGAACGGTGTTTATATTGCC<br>R: GTTCCTCCTTCCCACCCATACTGCTGCAATTTTATATACGTTC | pBSB2ctaA |
| *PnarG-mCherry* | F: CCGCGGGCTTTCCCAGCGGCAGTGTCCGTTTTATGGACAC<br>R: GTTCCTCCTTCCCACCCGAGTCAGGTGATGCTAAGTTCAC | pBSB8narG |
| *PskfA-mCherry* | F: CCGCGGGCTTTCCCAGCGCTGCCCTGCATCTCGGTTGTG<br>R: GTTCCTCCTTCCCACCAATTTTTGCATAGAGTCTATTGACATAG | pBSB8skfA |

| | | |
|---|---|---|
| *PcomGA-gfpmut3* | F: CCGCGGGCTTTCCCAGCTCCGATTACAGCTCTGGGTGCC<br><br>R: GTTCCTCCTTCCCACCCGCATATTGTAGAAAAAGAAGAAAAGG | pBSB2comGA |
| *PaprE- mCherry* | F: CCGCGGGCTTTCCCAGCCTGCTATCAAAATAACAGACTCGTG<br><br>R: GTTCCTCCTTCCCACCAATTCAGAGTAGACTTACTTAAAAGAC | pBSB8aprE |
| *PcggR- mCherry* | F: CCGCGGGCTTTCCCAGCCGCCTATACATTTTGGATCTTTGCGGTGATTAACAT<br><br>R: GTTCCTCCTTCCCACCCTTTTTTGCTGGACATTATATGTCCCGCTA | pBSB8cggR |
| *capE- mCherry* | F: CCGCGGGCTTTCCCAGCCAAGAGTATGACAATGATCCAAATG<br><br>R: GTTCCTCCTTCCCACCTGAATTATTTATTGGCGTTTACCGG | pBSB8capE |
| *PspoIIGA- mCherry* | F: CCGCGGGCTTTCCCAGCCGTTTACCATTCGTATGCCGCTGA<br><br>R: GTTCCTCCTTCCCACCCTTGCCTCACGCTGTTCCCCTTC | pBSB8spoIIGA |
| *PspoVC- mCherry* | F: CCGCGGGCTTTCCCAGCGAAGTTCCGATTCATCTGACCGGAG<br><br>R: GTTCCTCCTTCCCACCTCACATAACTCCCGTCTTCATAAAC | pBSB8spoVC |
| *PtapA- mCherry* | F: CCGCGGGCTTTCCCAGCGGTCCTTCAAAAAATGGAGGACC<br><br>R: GTTCCTCCTTCCCACCACACTGTAACTTGATATGACAATCG | pBSB8tapA |
